# A unified role for membrane-cortex detachment during cell protrusion initiation

**DOI:** 10.1101/696211

**Authors:** Erik S. Welf, Christopher E. Miles, Jaewon Huh, Meghan K. Driscoll, Tadamoto Isogai, Jungsik Noh, Andrew D. Weems, Joseph Chi, Theresa Pohlkamp, Kevin Dean, Reto Fiolka, Alex Mogilner, Gaudenz Danuser

**Affiliations:** University of Texas Southwestern Medical Center, Dallas TX, USA; New York University, New York NY, USA

## Abstract

Cell morphogenesis employs a diversity of membrane protrusions. They are discriminated by differences in force generation. Actin polymerization is the best studied mechanism of force generation, but growing interest in how variable molecular conditions and microenvironments alter morphogenesis has revealed other mechanisms, including intracellular pressure. Here, we show that local depletion of membrane cortex links is an essential step in the initiation of both pressure-based and actin-based protrusions. This observation challenges the quarter-century old Brownian ratchet model of actin-driven membrane protrusion, which requires an optimal balance of actin filament growth and membrane tethering. An updated model confirms membrane-filament detachment is necessary to activate the ratchet mechanism. These findings unify the regulation of different protrusion types, explaining how cells generate robust yet flexible strategies of morphogenesis.

## Main text

Cells use several mechanisms to generate the force required for membrane protrusion, with the choice of mechanism depending on cell-autonomous and environmental factors. High resolution microscopy of the molecular dynamics underlying these processes has until recently only been possible on cells adhering to glass coverslips. As a result, the most widely studied type of protrusions are the broad, flat lamellipodia that form in highly adherent cells *(1)*. Lamellipodial protrusions are driven by actin polymerization against the plasma membrane. The persistence of this process varies widely between cell types, from tens of seconds to minutes, dependent on the efficiency of regulatory processes that upregulate the rate of monomer incorporation into the filamentous actin (f-actin) network against the resistance of the plasma membrane and the coupling of the network to substrate-anchored adhesions *(2)*. In contrast, cells in more complex microenvironments often exhibit rounded protrusions, sometimes referred to as blebs, which are driven by intracellular pressure. Blebs form via separation of the plasma membrane from the actin cortex during a rapid expansion and typically persist for approximately 30 seconds before onset of a retraction phase that is characterized by recruitment of ezrin-radixin-moesin (ERM) family proteins and reassembly of an actin cortex *(3, 4)*. Perturbation of ERM proteins affects bleb size and frequency as well as bleb-driven migration *in vivo (5, 6)*, but this result is often ascribed to perturbations of the roles ERM proteins play in generating intracellular pressure *(7, 8)*. Other recently described protrusion mechanisms are hybrids between actin- and pressure driven mechanisms *(9–11).* The purely actin-driven lamellipodia and pressure-driven blebs thus represent the archetypical extremes of a wide protrusion spectrum. Especially in the context of cancer, the plasticity between lamellipodia-driven, mesenchymal migration and bleb-driven, amoeboid migration is a powerful mechanism for metastatic cells to navigate highly variable microenvironments *(12).* However, it has remained unclear how divergent the molecular processes underlying these protrusion types are and whether common pathways facilitate efficient interconversion in cells with high migration plasticity.

Metastatic melanoma cells in 3D microenvironments demonstrate morphological plasticity by switching between actin-rich protrusions and pressure-driven blebs. Therefore, they create an opportunity for the study of the two types of protrusions in the same cells. Using isotropic light sheet fluorescence imaging *(13, 14)* to monitor a GFP-tagged form of ezrin, we found that in agreement with previous reports *(15)* ezrin localized at the cell rear, while blebs formed at the front where the cell was burrowing into the collagen (Fig 1A, Movie 1). Projections of local membrane motion (Fig 1B, Movie 2) and ezrin concentration (Fig 1C, Movie 3) from 3D image data onto the cell surface showed that even outside of the high-ezrin uropod, protrusion is restricted to regions of lowest ezrin concentration. More detailed examination indicated that ezrin was reduced just as an individual bleb appeared, suggesting that ezrin is specifically depleted in these pressure-based protrusions (Fig 1D, Movie 4). Using geometrical classification via machine learning *(16)*, we identified individual blebs on the cell surface and quantified protrusive motion of these blebs as a function of ezrin intensity. This analysis revealed that bleb surfaces with the lowest ezrin intensity exhibited most forward motion, while bleb surfaces with the highest ezrin advance the least, supporting the hypothesis that ezrin depletion facilitates protrusion (Fig 1E).

**Fig. 1.**
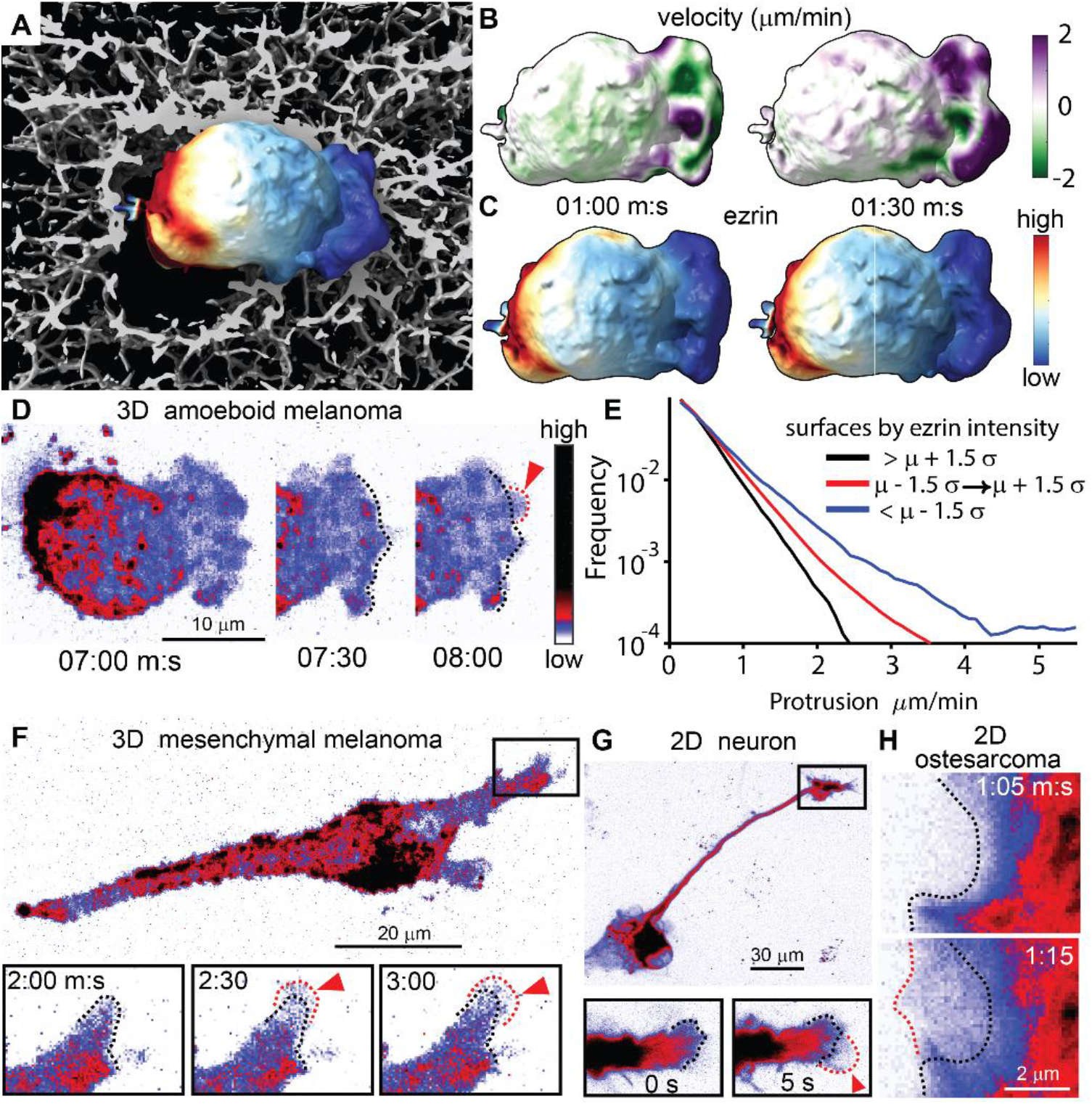
Diverse protrusive structures contain reduced ezrin. **(A)** Surface rendering of melanoma cell in 3D collagen with intensity of GFP-ezrin mapped to cell surface. Time-lapse data of the cell shown in (A) was used to quantify surface motion **(B)** and ezrin intensity **(C)** as a function of time. **(D)** Maximum intensity projection (MIP) of GFP-ezrin intensity shows detail of the time-lapse data from (A). **(E)** Quantification of surface motion on blebs as a function of ezrin intensity measured in 21 cells. Data is displayed as relative frequency in groups separated by ezrin intensity either above, below, or within +/-1.5 times the standard deviation of all measurements. **(F)** MIP of GFP-ezrin in a melanoma cell in 3D collagen. **(G)** rat cortical neuron. **(H)**. Time-lapse image sequence of GFP-ezrin in a MIP of GFP-ezrin in a primary U2OS osteosarcoma cell. Insets in (F) and (G) show magnification of protrusion time-lapse. In panels (D), (F), (G), and (H), black dashed lines show position of stable cortex and red dashed lines show nascent protrusion. Movies 1-6 show the temporal progression of these data.

We were surprised to observe that in the same melanoma cells, when they switch to mesenchymal migration, ezrin was depleted also from actin-driven protrusions (Fig 1F, Movie 5). We sought to confirm this observation in two other canonical forms of actin-driven protrusion, namely dynamic growth cones in rat cortical neurons (Fig 1G, Movie 6), and the flat lamellipodia of highly adherent osteosarcoma cells (Fig. 1H, Movie 7). Both protrusions also exhibited ezrin depletion (Fig 1G & H). Analysis of 3D light sheet microscopy data confirmed that this reduction in ezrin during protrusion was not due to protrusion thinning and thus size-related reduction of ezrin (Supplementary Figure 1). Together, these results suggested that ezrin depletion may be the unifying initiation event that drives both pressure driven and actin-driven protrusion.

While the role for membrane-cortex detachment in pressure-based protrusion is intuitive, our observation of ezrin depletion in actin-driven protrusions seems paradoxical because it has been proposed that ezrin localization stimulates actin polymerization *(17)*. To understand this conundrum we turned to the established theoretical model of actin-driven protrusion, which presumes that addition of actin monomers to the f-actin network pushes the cell membrane forward. First proposed in 1993, the Brownian Ratchet model explains how actin polymerization generates force *(18)*. Incorporating the effects of the angular distribution of actin filaments *(19)* and the effect of membrane tethering on force generation by actin *(20)* had improved the accuracy of the force-velocity relationship in actin-driven systems.

However, this model only explains how actin polymerization generates force once the edge is in forward motion, but cannot not explain protrusion initiation. The tethered Brownian Ratchet model requires membrane cortex attachment via WAVE and Arp2/3 to nucleate branched actin filaments but also requires membrane-cortex detachment to facilitate actin monomer addition to the ‘working’ filaments (Fig. 2A). Forces generated by the working filaments and those holding the membrane to the growing network are balanced and thus the model only applies to steady state membrane advancement. Given our observation that ezrin is reduced during actin-driven protrusion, we wondered if the membrane cortex attachment function of ezrin, which is distinct from that of WAVE, could initiate protrusion.

**Fig. 2.**
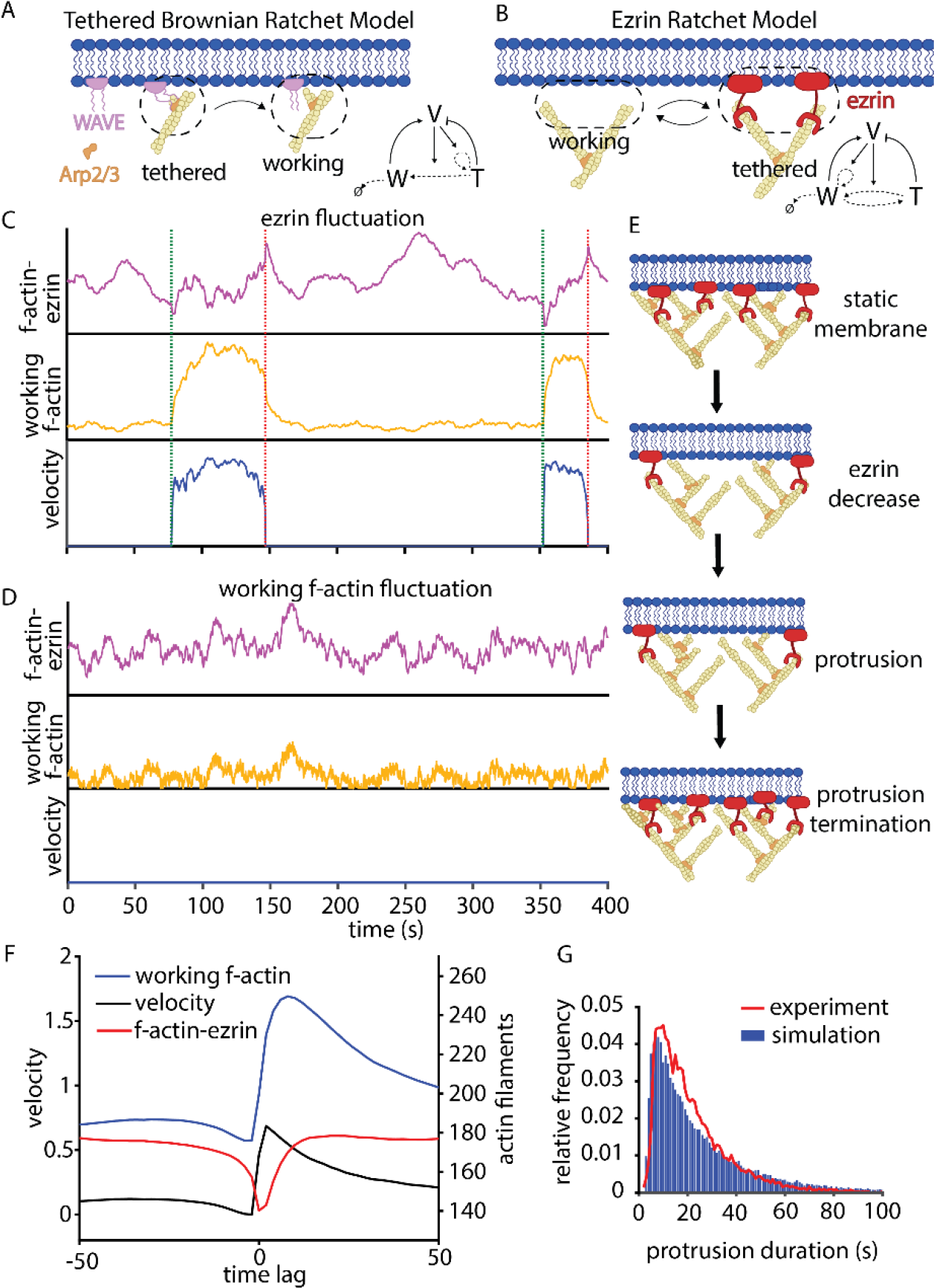
An ezrin ratchet model predicts that fluctuations in membrane-cortex attachment are required for protrusion initiation. **(A)** Illustration of the established tethered Brownian ratchet model. F-actin tethered to the membrane via WAVE is required to nucleate working actin filaments via arp2/3. Tethered filaments nucleate more tethered filaments but also restrict protrusion by coupling f-actin to the membrane. **(B)** Illustration of the tethered Brownian ratchet model accounting for the membrane cortex attachment function of ezrin. In contrast to the tethered Brownian ratchet model, working filaments can transition to tethered filaments but also self-nucleate because tethering is not exclusively coupled to WAVE. **(C&D)** Simulations of the ezrin ratchet model showing the effect of fluctuations either in ezrin concentration (C) or in actin filament density (D). In C&D, species concentrations are normalized and scaled the same for both conditions, and green and red dashed lines indicate protrusion onset and termination, respectively. **(E)** Illustration of the protrusion cycle according to the ezrin ratchet model: ezrin decreases, reducing membrane cortex attachment and allowing addition of actin monomers to push the membrane forward; protrusion terminates when ezrin binds to the increased actin filament concentration. **(F)** Mean value of total actin filaments and tethered filaments extracted from many simulated protrusion cycles, aligned to protrusion onset at 0 time lag. **(G)** Histograms of protrusion duration measured from experiments in U2OS cells or simulation of the ezrin ratchet model.

Upon adding the actin-membrane tethering function of ezrin to the Brownian ratchet model (Fig 2B), we found that ezrin fluctuations were sufficient to trigger protrusion initiation and that ezrin fluctuations coincided with protrusion initiation in the model simulation (Fig 2C). In contrast, fluctuations in the density of working actin filaments were not sufficient to initiate protrusion (Fig 2D). We explain this as follows: When ezrin levels are high enough to tether a critical fraction of filaments, then the protrusion stalls. Even if the density of actin filaments fluctuates up, the rapid on-off ezrin cycle simply increases the tethered filament subpopulations proportionally without affecting force balance and protrusion. However, if the ezrin level fluctuates down, the force balance shifts from tethered filaments (bound to ezrin) to working filaments, triggering protrusion (Fig 2E). Additional simulation results and the mathematical formalism of the Ezrin Ratchet Model are documented in the Supplementary Materials.

Analyzing the density of working actin filaments and ezrin-bound actin filaments in many simulated protrusion events predicts that the actin-bound form of ezrin decreases before protrusion starts (Fig 2F). The model also agrees with our previous experimental results showing that the actin filament density increases only after protrusion onset *(21)*. Further, the model suggests a mechanism for protrusion cessation, whereby the increase in actin filament density recruits ezrin to the cortex, increasing the filament – membrane tethers until the protrusion stalls. Remarkably, after adjusting the kinetics of free actin assembly and ezrin recruitment the Ezrin Ratchet Model quantitatively replicates the shape of the experimentally measured histogram of protrusion duration (Fig 2G). This suggests that the revised model captures the key relationships between protrusion initiation, propagation, and cessation.

To determine if experimental observations support the model predictions, we turned to adherent osteosarcoma cells, which exhibit repeated, localized actin-driven protrusions. Leveraging computational tools for localized sampling of the ezrin intensity at a constant distance to the moving cell edge and for statistical fluctuation analysis *(22)* we quantified the relationship between protrusion and ezrin concentration (Fig 3 A&B). In agreement with the predictions by the Ezrin Ratchet Model we found that the onset of ezrin depletion occurred prior to protrusion initiation (Fig. 3 C, D&E). Aggregating many such fluctuations over time and space (over 13,000 protrusion events from 1700 time series over 12 cells) allowed us to align ezrin intensity values with the quasi-stochastic velocity time series underlying spontaneous cell edge motion in order to determine the kinetics of ezrin depletion and recruitment during a stereotypical protrusion event. In support of the observations of single window time courses (Fig. 3C) and the predictions of the Ezrin Ratchet Model, these measurements confirmed that ezrin systematically decreases ~5s prior to protrusion onset (Fig 3 F). This relationship between ezrin and protrusion was strongest for ezrin time series sampled in the window layer subadjacent to the cell edge, where ezrin-mediated membrane-cortex tethers were expected to have highest density (Fig 3 G&H). The analysis also showed that ezrin concentration is at a minimum when protrusion velocity was at a maximum (Fig 3 I), and that it increased at the onset of retraction (Fig 3J). It reached a maximum ~3s after the peak in retraction velocity (Fig 3K). The observation that ezrin was recruited during retraction agreed with data showing that ezrin accumulated during bleb retraction *(3)*, suggesting that the molecular similarities between pressure-driven and actin-driven protrusions go beyond protrusion initiation. Notably, the ezrin species shown in the simulation results in Figure 2 is ezrin bound to actin. In contrast, the experimental data reflects both actin-bound and free ezrin. Therefore, we conjecture that the apparent increase in ezrin during retraction observed in the experimental data but not in the Ezrin Ratchet Model is due to aggregation of ezrin in the cytoplasm or in other unavailable stores near the membrane (Materials and Methods; Supplementary Figure 2).

**Fig. 3.**
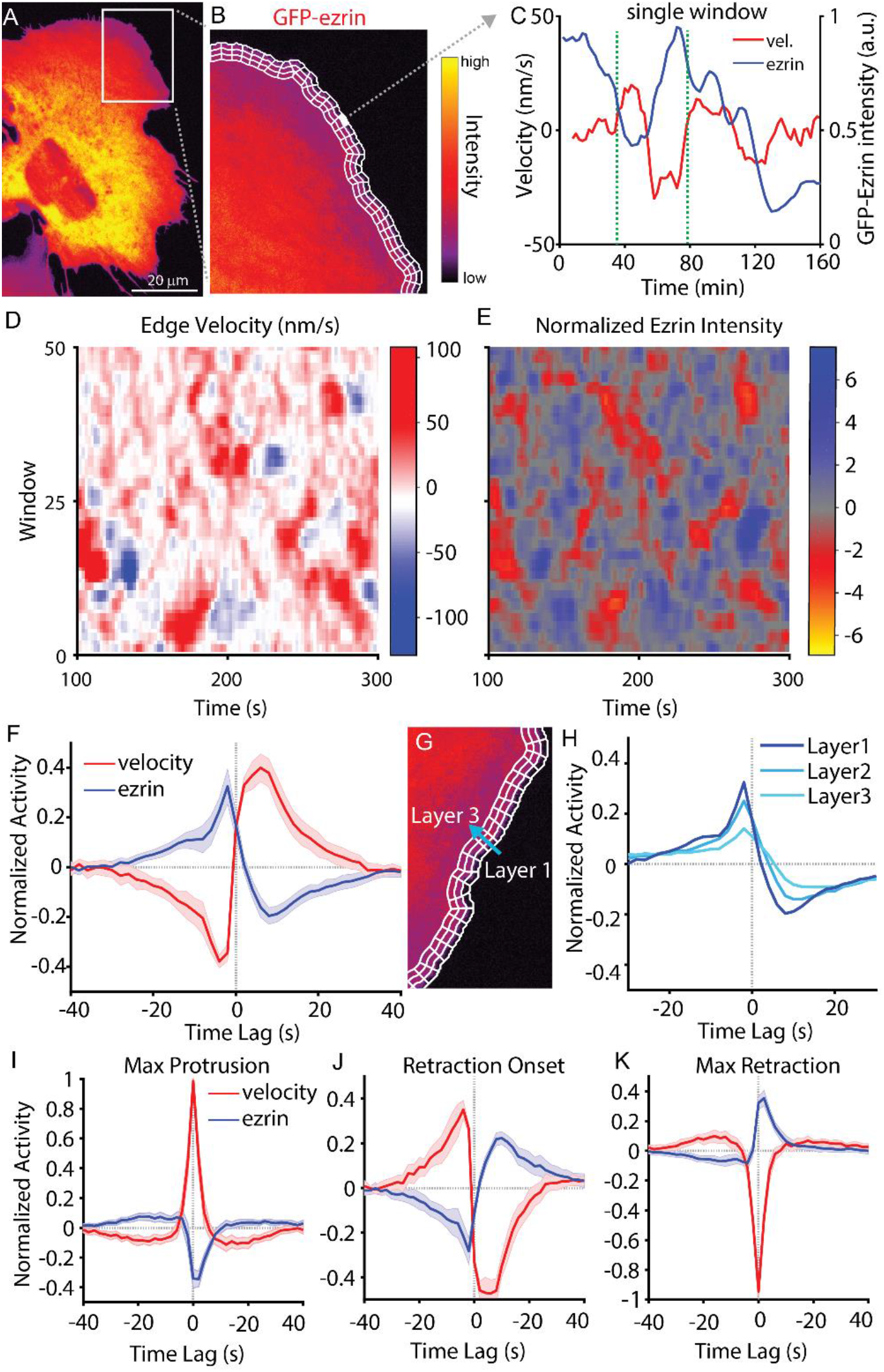
Stochastic local decreases in cortical ezrin initiate membrane protrusion. **(A)** Example image of U2OS osteosarcoma cell exhibiting lamellipodial protrusion. **(B)** Magnification of the lamellipodial region of the cell in (A) showing sampling windows used to extract time series of local protrusion and signaling. **(C)** Time series of ezrin concentration and cell edge velocity sampled in a single window as shown in panel (B). Green dashed lines indicate protrusion onset **(D)** Edge velocity map from a single movie of a U2OS cell exhibiting lamellipodial protrusion. **(E)** Ezrin intensity map for the movie in (D), with ezrin intensity normalized to exclude low frequency variation (Supp Fig 3). **(F)** Normalized activity of edge velocity and GFP-ezrin localization, aligned to protrusion onset in 12 U2OS cells imaged for 400 s each. Negative time lag indicates events before protrusion onset, whereas positive lag indicates events after protrusion onset. Shaded areas represent 95% confidence intervals about the mean activity. **(G)** Image of cell edge showing orientation of layers of sampling windows. (H) Normalized localization of GFP-ezrin in different sampling layers, showing the decrease in ezrin modulation with an increasing distance from cell edge. **(I, J, K)** Normalized activity of edge velocity and GFP-ezrin localization, aligned to (I) protrusion velocity maximum, (J) retraction onset, and (K) retraction maximum for the movies analyzed in (F). Line labels for I, J, and K are indicated in (I).

To further test the hypothesis that stochastic depletion of ezrin from the cortex initiates actin-driven protrusion, we exploited the biochemically well-characterized effect of ezrin phosphorylation on its affinity for cortical actin *(23)* (Fig 4A). A threonine to aspartic acid mutation (T567D) that mimics a phosphorylated form of ezrin with increased actin affinity causes impaired chemotaxis and migration *in vivo* and *in vitro (24, 25)*, but the defect this mutation introduces to the control of cell morphogenesis has remained unexplored. Expression of T567D ezrin resulted in dramatic morphological changes in both osteosarcoma cells cultured on 2D substrates and melanoma cells cultured in 3D (Fig 4B and Suppl. Figure 4). Likewise, increasing ezrin phosphorylation via knockdown of the ezrin-targeting phosphatase MYPT1 caused formation of smaller hemispherical protrusions (Suppl. Fig 5), whereas inhibition of ezrin function by the small molecule compound NSC668394 caused more and larger blebs to form (Supp Fig 6). Additionally, the recent finding that knockout of a different phosphatase with activity for ezrin decreases protrusion in neurons *in vivo* supports the role of ezrin dephosphorylation in protrusion initiation *(26)*. Compared to cells expressing wildtype ezrin, osteosarcoma cells expressing the T567D mutant exhibit fewer protrusions (Fig. 4B). Importantly, GFP-labeled T567D ezrin follows the same dissociation/reassociation pattern in the remaining protrusions as wildtype GFP-ezrin in control cells (Fig 4C). Thus, increased ezrin affinity for f-actin does not fully abrogate ezrin fluctuations, and these fluctuations remain permissive to the initiation of protrusion events. Similarly, we conjecture that the microvilli-like spikes in 3D melanoma harboring the T567D mutant represent stabilization of transient structures, such as filopodia, by increased cortical ezrin localization.

**Fig 4.**
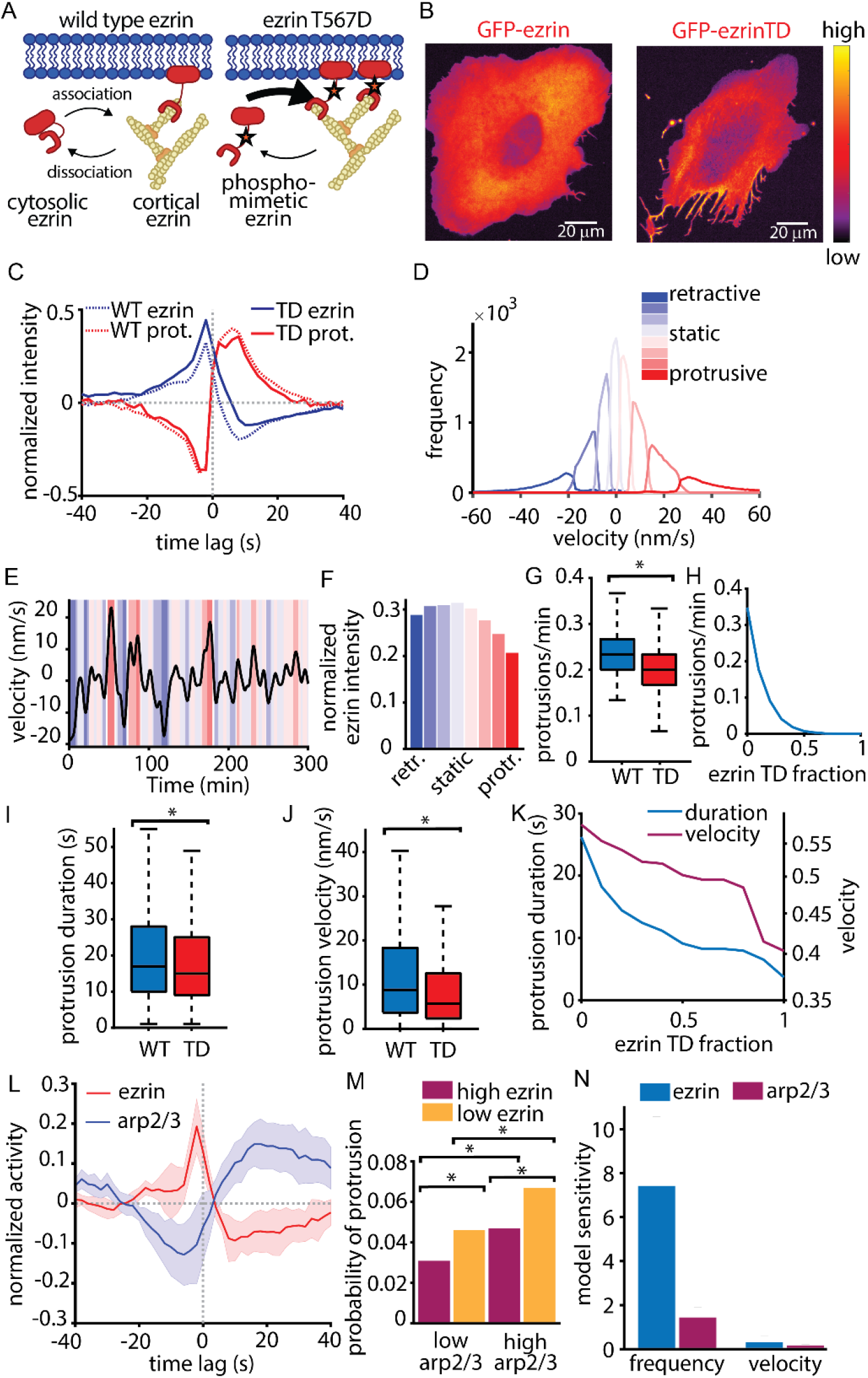
Ezrin fluctuations are regulated by phosphorylation and synergize with actin polymerization to control protrusion dynamics. **(A)** Illustration showing how the T567D (TD) mutation with higher affinity for actin increases membrane tethering compared to wild type (WT) ezrin. **(B)** Example images showing the effect of ezrin TD expression in the 2D osteoscarcoma model. **(C)** Normalized activity of edge velocity and GFP-ezrin localization, aligned to protrusion onset in 12 U2OS cells imaged in 12 cells for the WT condition and 9 cells for the TD condition. **(D)** Histogram showing how Hidden Markov Modeling (HMM) divides cell edge velocity according to edge velocity. **(E)** Example velocity time series from a single sampling window broken down into eight states classified by HMM. Colors defined in (D). **(F)** Normalized ezrin intensity during each of the eight states defined in (D). **(G)** Protrusion frequency in cells expressing either WT ezrin or TD ezrin, in addition to endogeneous ezrin, p = 0.039 via two sample t-test; n = 12 (WT) and n = 6 (TD). **(H)** Simulations of the ezrin ratchet model predicting the effect of increasing TD expression on protrusion initiation frequency. **(I)** Protrusion duration in cells expressing either WT ezrin or TD ezrin, p = 0.025 via two sample t-test comparing the ‘μ’-parameter of a Lognormal distribution fit to the protrusion durations in each cell (Suppl. Fig 8); n = 12 (WT) and n = 6 (TD). **(J)** Protrusion velocity in cells expressing either WT ezrin or TD ezrin, p = 0.027 via two sample t-test; n = 12 (WT) and n = 6 (TD). **(K)** Simulations of the ezrin ratchet model showing the effect of adding ezrin with increased actin affinity on protrusion duration and velocity. (L) Normalized activity of GFP-ezrin and HaloTag-arp2/3, aligned to protrusion onset (t = 0) in 8 U2OS cells. **(M)** Probability of protrusion in sampling windows categorized by HMM of ezrin and arp2/3 intensity. p < 0.01 for all comparisons via two sample t-test comparing probability per cell, n = 5 cells. **(N)** Mean sensitivity of model simulation results for protrusion frequency and velocity calculated over 10 different parameter values (Suppl. Fig 9).

To determine if expression of ezrin T567D alters the frequency of protrusion initiation events, we analyzed the edge velocity data with a hidden Markov model (HMM), which divides the velocity time series of each sampling window at the cell edge into a series of distinct motion states while eliminating spurious velocity fluctuations within the states (Fig. 4D&E). First applied to 1710 time series over 12 cells expressing WT GFP-ezrin, the HMM distinguished 4 protrusive and 4 retractive states. While the ezrin concentration varied only slightly between retractive states, it sharply decreased between protrusion states of increasing mean velocity (Fig. 4F). Next we identified the switches from any retractive to any protrusive state and compared their frequencies between cells expressing WT vs T567D ezrin (Suppl. Figure 7). Indeed, the higher affinity of T567D ezrin decreased the chances for a protrusion onset (Fig 4G). To determine if the Ezrin Ratchet Model recapitulates this result, we simulated the experimental expression of mutant ezrin by adding a second ezrin species with higher affinity for actin. We gradually increased the ratio of high to normal affinity ezrin and found a monotonically decreasing frequency of protrusion onsets (Fig 4H). We also observed a decrease in protrusion duration and protrusion velocity upon expression of mutant ezrin (Fig 4I&J), which was reproduced by simulation of the Ezrin Ratchet Model (Fig 4K). This result is consistent with previous data showing that increasing membrane tension after protrusion onset is rate-limiting for the speed and persistence of a protrusion event *(2)*. Elevated actin–membrane tethering by high-affinity ezrin would have a similar effect, leading to earlier termination of the event at lower velocities.

Our previous work on protrusion mechanics indicated that the growth of branched actin networks primarily serves the reinforcement after protrusion onset of actin assembly against mounting membrane tension *(21, 27)*. Therefore, we expected that the canonical nucleator of branched network formation, Arp2/3, would be recruited after protrusion onset and primarily after ezrin depletion. Indeed, concurrent imaging of Arp2/3 and ezrin in the same cell showed that Arp2/3 recruitment increased after protrusion onset and reached a maximum well after ezrin reached minimal levels (Fig 4L). Nonetheless, we were wondering whether recruitment of Arp2/3 still plays a role also in the protrusion onset. To address this we built a new HMM that identified intervals that were enriched vs depleted in ezrin or Arp2/3 and measured the frequency of protrusion within those intervals. As expected, intervals of low ezrin were enriched for protrusion compared to intervals of high ezrin; but intervals of low ezrin and high Arp2/3 exhibited a further increase in the probability of protrusion (Fig 4M), suggesting a synergistic effect of ezrin depletion and branched actin filament nucleation in protrusion initiation. To determine the relative importance of ezrin fluctuation or Arp2/3 activity for protrusion in the Ezrin Ratchet Model we performed simulations with progressively increasing values for the rate of branched actin polymerization, which reflects the activity of Arp2/3, and for the fraction of high affinity ezrin. The resulting changes in protrusion properties were normalized by the incremental change to the parameter value (Arp2/3 activity or ezrin affinity). Interestingly, this analysis predicts that ezrin has a larger influence on protrusion frequency than Arp2/3 and that ezrin and Arp2/3 have roughly equal influence on protrusion velocity (Fig 4N). Altogether, these results suggest that although branched actin polymerization aids protrusion initiation, its primary role is to reinforce growth of an actin network after a decrease in ezrin-mediated membrane cortex attachments.

The finding that fluctuations in ezrin concentration at the membrane initiate lamellipodia- and bleb-based protrusion raises the intriguing possibility that the trigger of protrusion events share common regulatory mechanisms regardless of the mode of force generation. In this case the mode of force generation determines the shape of protrusion, but not where and when it will occur. Clues that actin polymerization serves a protrusion reinforcement rather than initiation role abound: Quantitative fluorescent speckle microscopy analyses showed that the rate of actin assembly peaks after fastest protrusion *(2)*, the activity of actin polymerization factors Rac1, PI3K, and Arp2/3 all increase after protrusion onset *(21, 28, 29)*, and actin filament density increases when membrane tension increases *(30)*. Indeed, the positive feedback between protrusion and the transition between tethered and working actin filaments in the Ezrin Ratchet Model provides insight into how regulators of actin polymerization affect protrusion via amplification. This motif suggests that the canonical regulators of actin polymerization, such as soluble chemotactic or mechanical durotactic cues, amplify protrusion fluctuations but do not initiate them. This system design enables cells to randomly sample their environment via small edge fluctuations, facilitating identification of the path of least resistance in complex microenvironments *(31)*, while the commitment to directed, persistent protrusion is made later via the reinforcement module that ultimately guides directionality *(29, 32, 33)*. Altogether, these experimental and theoretical insights fundamentally change our understanding of cell morphogenesis, not only because they identify the elusive initiator of protrusion but because they unify seemingly different forms of protrusion under one regulatory mechanism.

## Author contributions

According to the CRediT system, author contributions are as follows. ESW: Conceptualization, Formal analysis, Investigation, Methodology, Project administration, Visualization, and original draft Writing. CJM: Conceptualization, Investigation, Methodology, and original draft Writing. JH: Formal analysis, Investigation, Methodology, Software. MKD: Conceptualization, Formal analysis, Investigation, Methodology, and Visualization. TI: Investigation. JN: Formal analysis, Investigation, Methodology, Software. ADW: Investigation, and Visualization. JC: Resources. TP: Resources. KD: Resources. RF: Methodology, Supervision, and Resources. AM: Conceptualization, Methodology, and original draft Writing. GD: Conceptualization, Funding acquisition, Methodology, Resources, and original draft Writing.

## Funding

This work was funded by CPRIT (RR160057 to RF), the National Institute of Health (K25CA204526 to ESW, F32GM116370 and K99GM123221 to MKD, R33CA235254 to RF, and R01GM071868 to GD), and the US Army Research Office W911NF-17-1-0417 to AM. ADW is a fellow of the Jane Coffin Child’s Memorial Fund.

## Competing Interests

The authors declare no competing interests.

## Supplementary Materials

Materials and Methods, including References (34–51)

Supplementary Figures 1-11

Movies 1-8

## Supplementary Materials

### Materials and Methods

#### Cell culture

MV3 cells were obtained from Peter Friedl (MD Anderson Cancer Center, Houston TX). U2OS cells were obtained from R. McIntosh (University of Colorado, Boulder CO). MV3 cells were cultured in DMEM (Gibco) supplemented with 10% fetal bovine serum (FBS; ThermoFisher) and U2OS cells were cultured in McCoy’s Medium (Gibco) supplemented with 10% FBS. The GFP-ezrin constructs were acquired from Addgene (plasmids #20680 and #20681; Hao JJ, Liu Y, Kruhlak M, Debell KE, Rellahan BL, Shaw S, J Cell Biol. 2009) and cloned into the pLVX-puro vector (Clontech). The Td-Tomato-membrane construct consists of the first 60 base pairs of GAP43 (neuromodulin) fused to Td-Tomato and inserted into the pLVX-neo vector (Clontech). Lentivirus was produced and cells were infected according to the manufacturer’s instructions (Clontech). Cell lines expressing lentiviral constructs were purified using either purimycin or neomycin (ThermoFisher). Cells containing knockdown of protein phosphatase 1 (regulatory subunit 12A) were created using the lentiviral pLKO.1 vector containing MISSION shRNA sequences (Sigma) per the manufacturer’s instructions. MV3 cells were treated with the Ezrin inhibitor NSC668394 using a 5 mM stock solution in DMSO. Final working concentration was 10 uM NSC668394 (EMD Millipore) and 0.2% DMSO (v/v) in phenol red free DMEM w/ 10% FBS. Inhibitor concentration was chosen based on previously published in vitro protocols (PMID: 21706056). MV3 cells expressing GFP-F-tractin *(34)* (Yi, J., Wu, X.S., Crites, T., and Hammer, J.A. Mol. Biol. Cell, 2012) were mounted in bovine collagen as previously described. Cells were imaged immediately before drug treatment, and then again 20 minutes after treatment.

#### 3D sample preparation

Collagen gels were created by mixing bovine collagen I (Advanced Biomatrix) with concentrated phosphate buffered saline (PBS) and water for a final concentration of 2 mg/mL collagen. This collagen solution was then brought to pH 7 with 1N NaOH and mixed with cells just prior to incubation at 37°C to induce collagen polymerization. Cells were suspended using trypsin/EDTA (Gibco), centrifuged to remove media, and then mixed with collagen just prior to incubation at 37°C to initiate collagen polymerization. To image collagen fibers, a small amount of collagen was conjugated directly to AlexaFluor 568 dye and mixed with the collagen sample just prior to polymerization.

#### 3D cell imaging

3D samples were imaged using either an axially scanned light sheet microscope *(14, 35)* or using our meSPIM microscope *(13)*, both of which provide nearly isotropic, diffraction-limited 3D images. Samples were imaged in phenol red free DMEM containing 25mM HEPES (ThermoFisher) with 10% FBS and antibiotic-antimycotic (Gibco), held at 37°C during imaging. For cells on coverslips imaged in 3D, the focused area of the light sheet was scanned parallel to the coverslip in order to increase imaging speed without sacrificing resolution *(35)*. Images were collected using sCMOS cameras (Orca Flash4.0 v2, Hamamatsu) and microscopes were operated using custom Labview software. All software was developed using a 64-bit version of LabView 2016 equipped with the LabView Run-Time Engine, Vision Development Module, Vision Run-Time Module and all appropriate device drivers, including NI-RIO Drivers (National Instruments). Software communicated with the camera via the DCAM-API for the Active Silicon Firebird frame-grabber and delivered a series of deterministic TTL triggers with a field programmable gate array (PCIe 7852R, National Instruments). These triggers included analog outputs for control of mirror galvanometers, piezoelectric actuators, laser modulation and blanking, camera fire and external trigger. All images were saved in the OME-TIFF format. Some of the core functions and routines in the microscope control software are licensed under a material transfer agreement from Howard Hughes Medical Institute, Janelia Farm Research Campus.

#### 3D image rendering and analysis

3D image data was processed as described previously *(13, 16)*. Briefly, image data was segmented to create a surface represented as a 3D triangle mesh, using either a manually selected intensity threshold or using Ilastik *(36)*. The triangle meshes shown in Figure 1 A, B, and C were rendered in ChimeraX *(37)*. Colored triangle meshes were exported from Matlab as Collada.dae files using custom-written code and were rendered using full lighting mode. To measure the fluorescence intensity local to each mesh face, we used the raw, non-deconvolved, fluorescence image. At each mesh face, we used a kd-tree to measure the average pixel intensity within the cell and within a sampling radius of the mesh face. To correct for surface curvature dependent artifacts, we depth normalized^33^ the image prior to measuring intensity localization by normalizing each pixel by the average pixel intensity at that distance interior to the cell surface. Prior to analysis, we also normalized each cell’s surface intensity localization to a mean of one.

#### 2D imaging

U2OS cells imaged in 2D were plated on glass coverslips coated with 10 mg/mL fibronectin and imaged on a Nikon EclipseTi-E inverted motorized microscope coupled to an Andor Diskovery TIRF/ Borealis widefield illuminator equipped with an additional 1.8x tube lens (yielding a final magnification of 108x). The microscope was equipped with a 60x Nikon 1.49 NA TIRF DIC objective, Andor Zyla 4.2 16 bit, 100 fps, 2048×2048 px sCMOS cameras, and OKO lab custom built full body environmental chamber with temperature control and CO2 stage incubator.

#### Analysis of temporal relationships between protrusion and GFP-ezrin intensity

We performed several experiments to confirm that our observation of ezrin depletion before protrusion was not an artifact. To determine if the newly protruding region of lamellipodium is simply thinner than the established cortical region, we performed 3D light sheet microscopy of cells adhering to glass coverslips. Optical reslicing of this 3D data to show the profile of a cell during lamellipodial protrusion confirms that, within the limitation of the diffraction limit of light imaging, the newly protruding lamellipodium does not appear to be substantially thinner than the established lamellipodium (Supplementary Figure 1). To confirm this result quantitatively and to exclude the possibility that a lamellipodial protrusion somehow restricts access of fluorescent markers from entering a protrusion, we analyzed the relationship between protrusion and intensity of a freely diffusing fluorophore in the cell’s cytosol. This analysis shows that the intensity of a cytosolic fluorophore in the second layer shows no relationship with protrusion onset or maximum velocity (Supplementary Figure 10). Thus, we conclude that the reduction in ezrin is not due to a thinner protrusion or restricted access of a fluorophore to a protrusion. Finally, to eliminate any inaccuracies in the identification of the cell’s edge during protrusion, we performed image segmentation and edge velocity measurement on images of a cell membrane marker (tD-Tomato membrane) instead of GFP-ezrin. Linescans of a protrusion clearly show the cell edge movement via the td-Tomato membrane signal, whereas the GFP-ezrin signal is reduced in the protrusion, although it quickly reaches similar intensity to the non-motile areas after protrusion (Supplementary Figure 11). The GFP-ezrin intensity measurements were aligned to windows determined based on image masks of the tD-tomato membrane images, so GFP-ezrin intensity measurements at the cell edge were not biased by the concentration of GFP-ezrin.

#### Time series analysis

To study dynamic subcellular activities relative to edge motion, we computationally tracked the cell boundary movement over time and subsequently defined a cell-shape invariant coordinate system allowing registration of movement and signaling. The cell boundaries were segmented using intensity thresholding using the Td-Tomato-membrane so that possible reductions in GFP-ezrin at the cell edge would not result in an inaccurate cell edge measurement. To calculate locally the displacement of the cell edge we morphed the segmented cell outlines between consecutive time points using the morphodynamic profiling algorithm previously described *(38)*.

Upon definition of the cell edge motion the segmented cell masks were partitioned into sampling windows of size 8×4 pixels (~1 μm x 0.5 μm) using contour lines and ridges in the Euclidean distance transform map to the cell edge. One of the windows within the outermost layer at the first time point was set to be the origin. The location was propagated through time frames using the information of edge displacements calculated as above.

To identify the edge motion events, we first smoothed the edge velocity map by representing the motion time series of an individual window by a smoothing spline, computed with a Matlab function csaps() and manually chosen smoothing parameters. We identified the time points and locations where the smoothed velocities are positive (negative) as a protrusion (retraction) phases. Protrusion/retraction periods shorter than 25 seconds were excluded. Within each sampling window, the beginning time points of protrusion/retraction phases was then detected as the protrusion/retraction onsets. Within each protrusion/retraction period, the time points with the maximum/minimum smoothed velocities were also detected. To capture the local dynamics of ezrin intensity around the edge motion events, we first normalized time courses of ezrin intensity in each window by subtracting its mean and dividing by its standard deviation. We then locally sampled the normalized activities within ±40 sec before and after the motion events. Additional information about the computation was previously described (Azoitei et al. 2019. JCB, under revision).

#### Hidden Markov modeling

To identify the temporal behaviors of edge velocity maps, we implement the hidden Markov modeling to the velocity map data, computed with the R package ‘depmixS4’ and the functions within (39). The modeling works with an input of pre-determined number of states and time series data which output the hidden state time series determined by the average and variance of the data within each state. The seed is initialized prior to model fitting by the function set.seed() for consistent output.

To have uniform criteria of state selection among different cells, we concatenate all the velocity time series into a single long vector with missing values connected. The data is saved as a depmix object and computed using the fit() function without setting initial transition probability. The package also supports multivariate time series computation that seems appropriate for our data. However, velocity data that is outside the range of 5 standard deviations is imputed as a NaN (Not a number) which give us time series with missing data. To deal with missing values within windows, we decided to consider the whole data as a single vector. The model computation is iterated until there is no significant change in likelihood.

The HMM was computed with increasing number of states until the minimum proportion of a single state reached 5 %. For the case of 12 WT cells and 6 TD cells, 8 states were chosen which gave good interpretation of the states by their average velocity (Figure 4D). This method provides an interpretable clustering of time series data superior to Gaussian mixture modeling (GMM). The advantage of the HMM over the GMM is that the state isn’t strictly determined by a single time point but its previous data point. This gives a much smoother state selection that well represents the temporal dynamics of edge velocity maps.

For the temporal behaviors of Ezrin & Arp2/3 localization, the Markov state selection was dominated by low frequency signals mostly representing the subcellular intensity level. Given that our interest was the sudden increase or decrease of signals related to the velocity dynamics, we implemented a low frequency normalization technique that adjusts the width of the temporal autocorrelation of the Ezrin intensity signal to the width of the temporal autocorrelation of the velocity fluctuations (Supplementary Figure 3). Using the fact that the full cycle of edge velocity is 40 frames (80 seconds), we subtract the median time series of a 60 frame moving window from the raw Ezrin time series. This method removes variations in the signals with a quasi-periodicity longer than 120 seconds, which are unrelated to variations associated with the protrusion/retraction cycles. After this low frequency normalization, the hidden Markov model was computed for the concatenated time series from 5 cells. The signals were classified into 4 states ordered by the average intensity levels. We further combine the top 2 and bottom 2 states as high and low activity states. This was due to the fact that the HMM modeling with 2 states were mainly driven by the local variance of the activities not the average intensity. This problem disappeared after choosing more than 3 states. To account for differential ezrin intensity values in different cells, the GFP-ezrin intensity for each cell was normalized to fall between 0 and 1. The values shown in Figure 4E represent the mean values over all windows in all cells measured.

#### Statistical comparisons

In our experience, the greatest source of variation in cell edge fluctuations arises from cell-cell variability. Therefore, statistical comparisons between cells expressing wild type and mutant ezrin are performed on a per cell basis. In order to show the spread of all data points, data shown in the box plots in Figure 4 is pooled from all cells.

Protrusion initiation frequency was analyzed by counting the total protrusion events per cell as classified by HMM velocity states with mean greater than zero. Events were counted per window and then a mean (protrusions/window) was calculated for each cell. To avoid influence of frequent spurious events, only protrusions longer than 25 seconds were included in this analysis. We were not able to identify an appropriate probability distribution function to model the single cell distribution of protrusion initiation events per window, so we used a two sample t-test with equal variances to compare protrusion initiation event frequency between cells expressing WT and TD ezrin on a per cell basis. A Wilcoxan rank sum test of this comparison yields a significance of p = 0.0182.

Protrusion duration was analyzed by fitting the protrusion durations, as determined by the HMM, in each cell to a Lognormal distribution and performing a two sample t-test with equal variances on the mu parameter (measure of the distribution’s central value) on the per-cell Lognormal distributions. A t-test of the medians of the distributions of durations from single cells expressing either ezrin WT or TD also showed significance (p = 0.048). A Wilcoxan rank sum test of the Mu parameter yields a significance of p = 0.0668. Because we were able to identify the Lognormal distribution as an appropriate model for protrusion duration, all protrusions, regardless of duration, were included in this analysis.

A probability distribution function that fits the velocity data for each cell could not be found, therefore cells were compared via a t-test of the mean protrusion velocities measured for each cell. These differences were also significant according to a Wilcoxan rank sum test (p = 0.0245). To exclude the effect of rare but numerically more influential protrusion velocity values, we excluded velocity values greater than 20 nm/s from this analysis. The excluded velocity measurements accounted for less than 20% of all measurements across all cells. All values are included in the box and whiskers plot in Figure 4I.

For the comparisons between Arp2/3 and ezrin levels shown in Figure 4M, the statistical tests were again performed on a cell-by-cell basis. For each condition, the Poisson ratio of protrusion events/total measurements windows was calculated per cell. T-tests on the resulting mean values produced p-values <0.01 for all comparisons shown in Figure 4M. A Wilcoxan rank sum test for these same comparisons produced p-values < 0.05.

#### Mathematical model and simulation

In this section of supplement, we provide details on the basic ezrin-ratchet model, as well as variations referenced in the main text.

#### Spatial Model

**Methods Figure 1.**
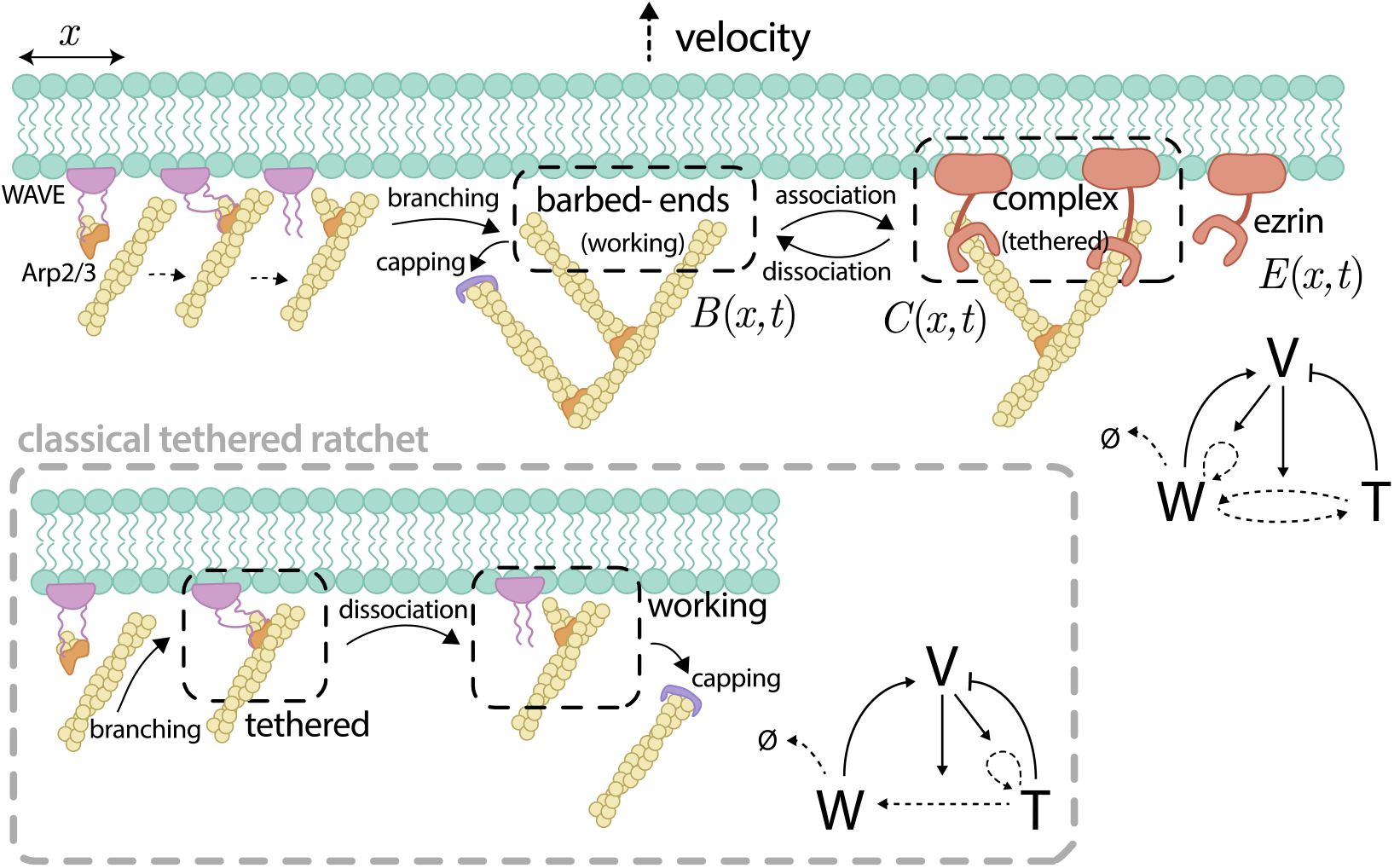
Cartoon schematic of the proposed mathematical model, contrasted with the classical tethered ratchet model (20).

The mathematical model of the ezrin ratchet system is based on the classical tethered ratchet model (20) and is graphically summarized in **Methods Figure 1**. The model describes the dynamics of three population densities along the leading edge of the cell: i) F-actin barbed ends, which we term working filaments, exert a force on the membrane through a polymerization ratchet (40), ii) ezrin-barbed end complexes, which we term actin-ezrin, serve as tether forces on the membrane, and iii) free ezrin, which binds to F-actin to form the complex. These three state variables, along with order-of-magnitude estimates of their values, are summarized in **Methods Table 1**. The estimated values are taken from previous modeling and experimental works *(41–43)*. The spatial component of these densities corresponds to the thin strip along the leading edge of the cell described by the coordinate 0 < *x* < *L*.

**Methods Table 1.**
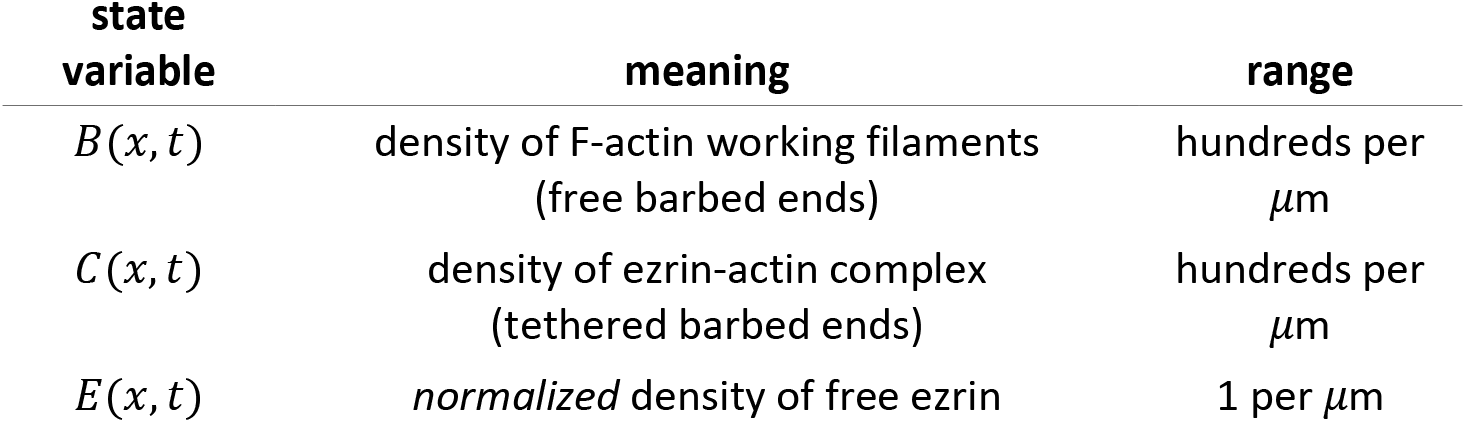
State variables of the spatial system.

| state variable | meaning | range |
| --- | --- | --- |
| $B(x,t)$ | density of F-actin working filaments (free barbed ends) | hundreds per $\mu\text{m}$ |
| $C(x,t)$ | density of ezrin-actin complex (tethered barbed ends) | hundreds per $\mu\text{m}$ |
| $E(x,t)$ | normalized density of free ezrin | 1 per $\mu\text{m}$ |

From **Methods Figure 1**, we note a key structural difference: in the classical tethered ratchet, new filaments enter the system as tethered. However, in this work, we assume this pool of tethered filaments is negligible compared to those that become transiently tethered after entering the system. Based on this, the dynamics of the model consist of first order binding and unbinding reactions described in (1).

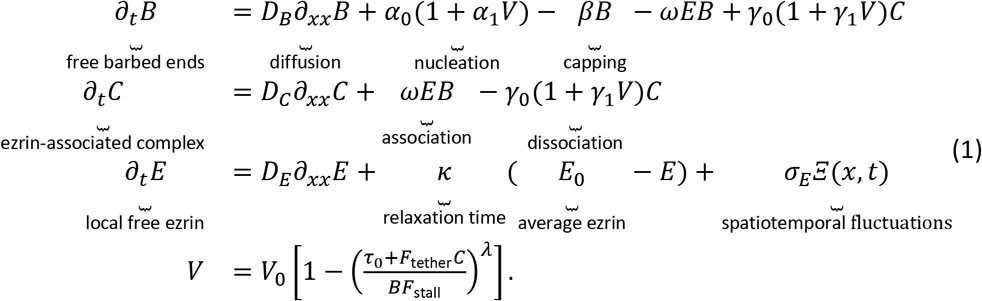

- The return of free ezrin into the pool of *E* is neglected, as we are assuming there is not a limiting supply, but rather a large number that fluctuates slightly in space and time. The mean value of free ezrin is also normalized to *E*_0_ = 1, as the effective binding rate *ωE* is the only quantity of interest, not the free ezrin level directly.
- The velocity of the membrane depends on the force-per-barbed end [6] and takes the functional form from empirical data *(43, 44)*.
- The branching rate increases with velocity *(43, 45)* due to geometric and force effects, which are included in the simplest way possible here.
- The disassociation rate between barbed ends and ezrin is also known to be force-dependent *(46)*.
- The model neglects membrane retractions and instead has a minimum velocity of zero. However, the force-velocity curve could be modified easily to include negative velocities for appropriate forces without meaningfully changing the results of the model.

Unless noted otherwise, the parameters used in all simulations are described in **Methods Table 2**.

**Methods Table 2.** Parameter values used for the mathematical model. The dagger symbol (†) denotes parameters used in the nonspatial version of the model.

| parameter | Meaning | range/source | value used |
| --- | --- | --- | --- |
| $D_B$ | barbed end diffusivity | $\sim v^2 \beta$ tenths of $\mu\text{m}^2/\text{s}$ (41) | .025 [$\mu\text{m}^2/\text{s}$] |
| $\alpha_0$ | barbed end branching rate | tens to hundreds per $\mu\text{m}$ per second (41) | 50 [$1/(\mu\text{m} \cdot \text{s})$] |
| $\alpha_1$ | branching rate increase from velocity | 0.5 to 2, estimated from data in (30, 44) | 1 |
| $\beta$ | capping rate | tenths per second (41) | .3 [$1/\text{s}$] |
| $\omega$ | barbed-end ezrin association rate | seconds (23) | 2.1 [$1/\text{s}$] |
| $\gamma_0$ | barbed-end ezrin dissociation rate | seconds (23) | 2 [$1/\text{s}$] |
| $\gamma_1$ | barbed-end ezrin dissociation rate increase | 0.5 to 2, estimated from data in (46) | 0.5 |
| $D_C$ | barbed-end/ezrin complex diffusivity | slow (23) | $10^{-4} [\mu\text{m}^2/\text{s}]$ |
| $D_E$ | free ezrin diffusivity | hundreths (23) to tenths (47) of $\mu\text{m}^2$ | $0.1 [\mu\text{m}^2/\text{s}]$ |
| $\kappa$ | relaxation time of ezrin fluctuations | estimated from data ∼ tenths of seconds | $0.025 [1/\text{s}]$ |
| $\sigma_E$ | magnitude of ezrin fluctuations | estimated from data $\sim 10\%$ | .045, .03<sup>†</sup> |
| $E_0$ | mean level of ezrin | normalized | $1 [1/\mu\text{m}]$ |
| $\tau_0$ | cortical tension | tens to hundreds of $pN$ per $\mu\text{m}$ (20, 41, 43, 44) | $25 [pN/\mu\text{m}]$ |
| $F_0$ | stall force in ratio of tethered/barbed ends | ones (20, 41) | 1 |
| $\lambda$ | steepness of force-velocity curve | $\sim 4 - 8$ (48) | 4 |
| $F_{\text{tether}}$ | tether force | piconewtons, estimated (20) | $1 [pN]$ |
| $L$ | domain size | experimental observations | $25 [\mu\text{m}]$ |

- The values of *ξ, σ_E_* are approximated from experimental measurements in this work (not shown). Specifically, the correlation time of ezrin fluctuations was seen to be 1/*κ* ≈ 40[s] and standard deviation approximately 10% of the baseline value, which is reproduced with these values. The diffusion in the spatial model dissipates fluctuations, so the value taken in this model is slightly higher than that taken in the nonspatial model, as noted in the table.
- The tether force *F*_tether_ reported in previous works (20) is an order of magnitude higher (tens of piconewtons), but we note that this force is proportional to the velocity of the membrane. Consequently, in this work, where the velocity is an order of magnitude slower, we expect tether forces on the magnitude of piconewtons.

The system of spatial PDEs (1) was simulated using a semi-implict Euler-Maruyama scheme *(49)* with timestep *dt* = *10e-3* until *tmax = 800* and 301 spatial grid points. A typical trajectory of the spatial model can be seen in **Methods Figure 2**.

**Methods Figure 2.**
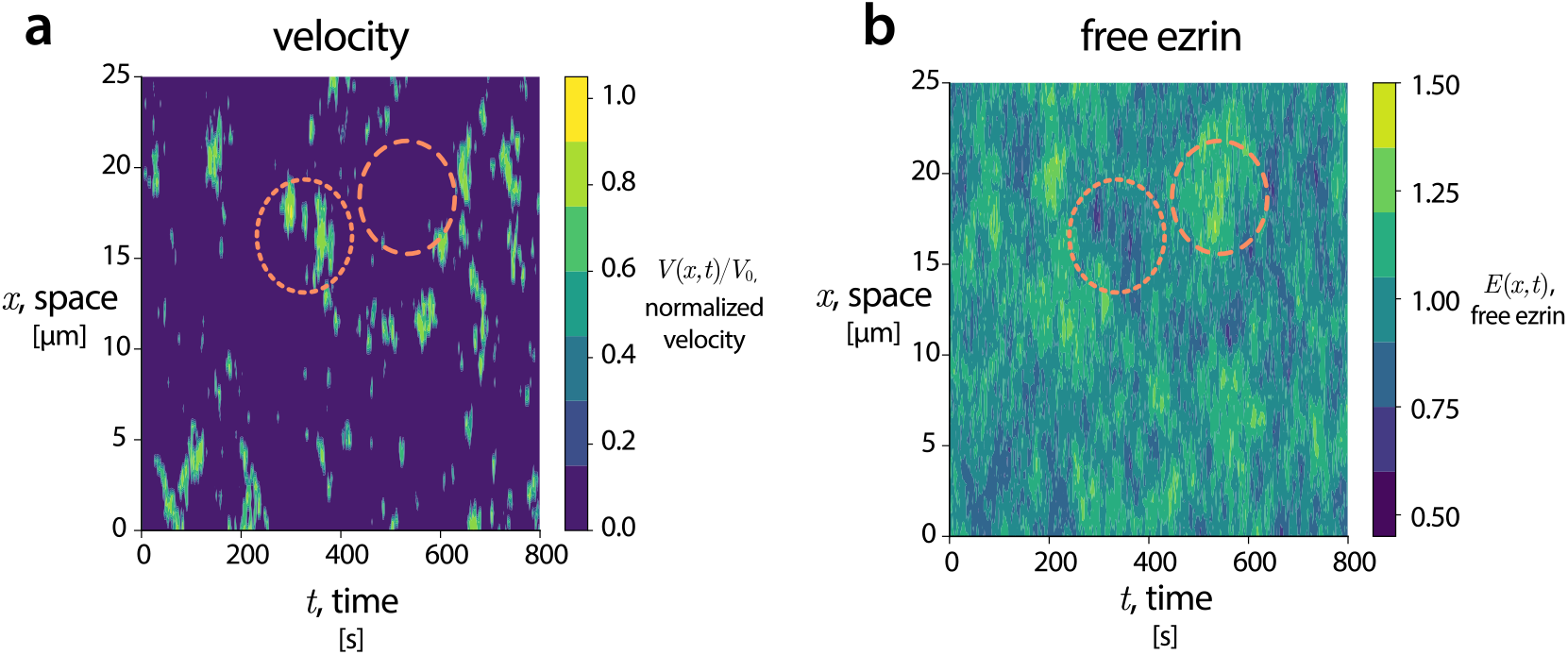
Typical simulation of the spatial system, (1). On the left, the instantaneous velocity determined by the relative quantities of free and tethered filaments. On the right, the density of free ezrin. The two highlighted regions qualitatively demonstrate that high protrusive activity corresponds to low ezrin free levels (dotted) and low protrusive activity corresponds to high ezrin levels (dashed).

From **Methods Figure 2**, we see that the mathematical model demonstrates transient protrusive behavior akin to the metastable switching of the canonical stochastic Allen-Cahn equation (50). From these simulations, we also note that protrusive activity correlates with low free ezrin levels. To understand this relationship more quantitatively, we turn to a simpler (nonspatial) model.

#### Nonspatial model

Spatial variations occur at a spatial scale that coincides with the windows of the experimental setup (microns) due to the relatively small diffusivities of the quantities in the system. Consequently, for ease of analysis, we instead study a non-spatial system with the same state variables as the spatial model, with the interpretation that these quantities are constant in each experimental spatial window. The nonspatial model is then

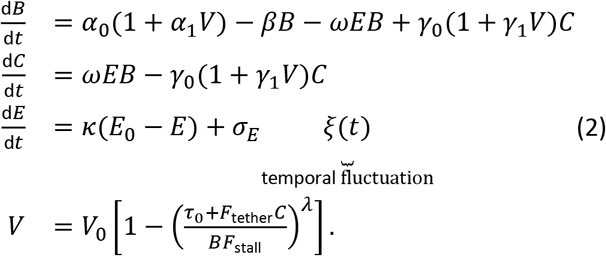

Statistics of protrusions in the experimental setup are computed on a window-by-window basis, and therefore are inherently nonspatial. Therefore, the main text figures, all statistics of the model (e.g. protrusion durations) are describing those obtained from the nonspatial system (2).

#### Stability & equilibria analysis

Although determining the equilibria (and stability thereof) of (2) is feasible, it becomes considerably less wieldy in the limit that *λ* → ∞, so the force-velocity curve becomes a sharp threshold. For the remainder of this subsection, we make that assumption.

In this limit, the force velocity curve becomes

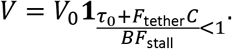

where **1**_(·)_ is 1 whenever the condition (·) is true and zero otherwise. Consequently, the behavior of the model can be split into two regimes:

**Regime 1:** 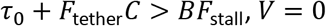.

**Regime 2:**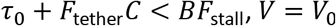.

We now derive the conditions such that each of these regimes provides a basin of attraction for a stable steady state, separated by the separatrix *τ*_0_ + *F*_tether_*C* = *BF*_stall_. In both regimes, *E*(*t*) → *E*_0_ stably.

**Regime 1.** Taking *E*(*t*) = *E*_0_ and *V* = 0, The system (2) becomes

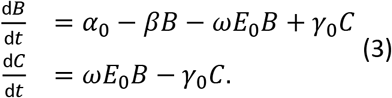

The system (3) has an equilibrium

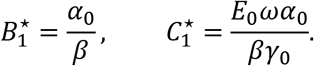

**Regime 2**. Taking *E*(*t*) = *E*_0_ and *V* = *V*_0_, The system (2) then becomes

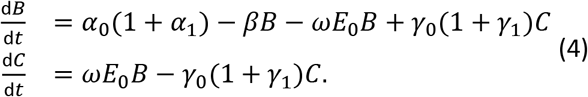

The system (4) has an equilibrium

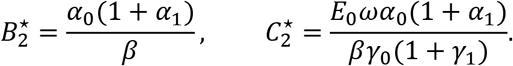

Thus, the conditions for these to both be stable correspond to them appearing in each of the appropriate regimes, so for region 1, substituting the equilibrium to the inequality condition yields

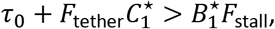

which provides the condition

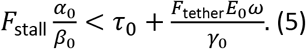

The condition (5) effectively says that a stall state exists if the steady-state force per working filament is smaller than the combination of membrane tension and tethered filaments. Similarly, in regime 2, we have

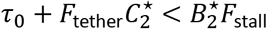

yielding

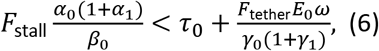

which has the same interpretation: during a protrusive event, the force per filament must be lowered due to the increased detachment rate of ezrin from filaments. Combining the conditions (5) and (6) yields the simultaneous condition

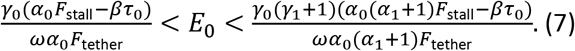

Although the functional form of (7) is complex, the lesson is intuitive: to drive transient protrusive events with this model, the baseline ezrin level must be at an intermediate sweet-spot where too little ezrin would have constant velocity but too much would have no protrusive activity at all.

**Methods Figure 3.**
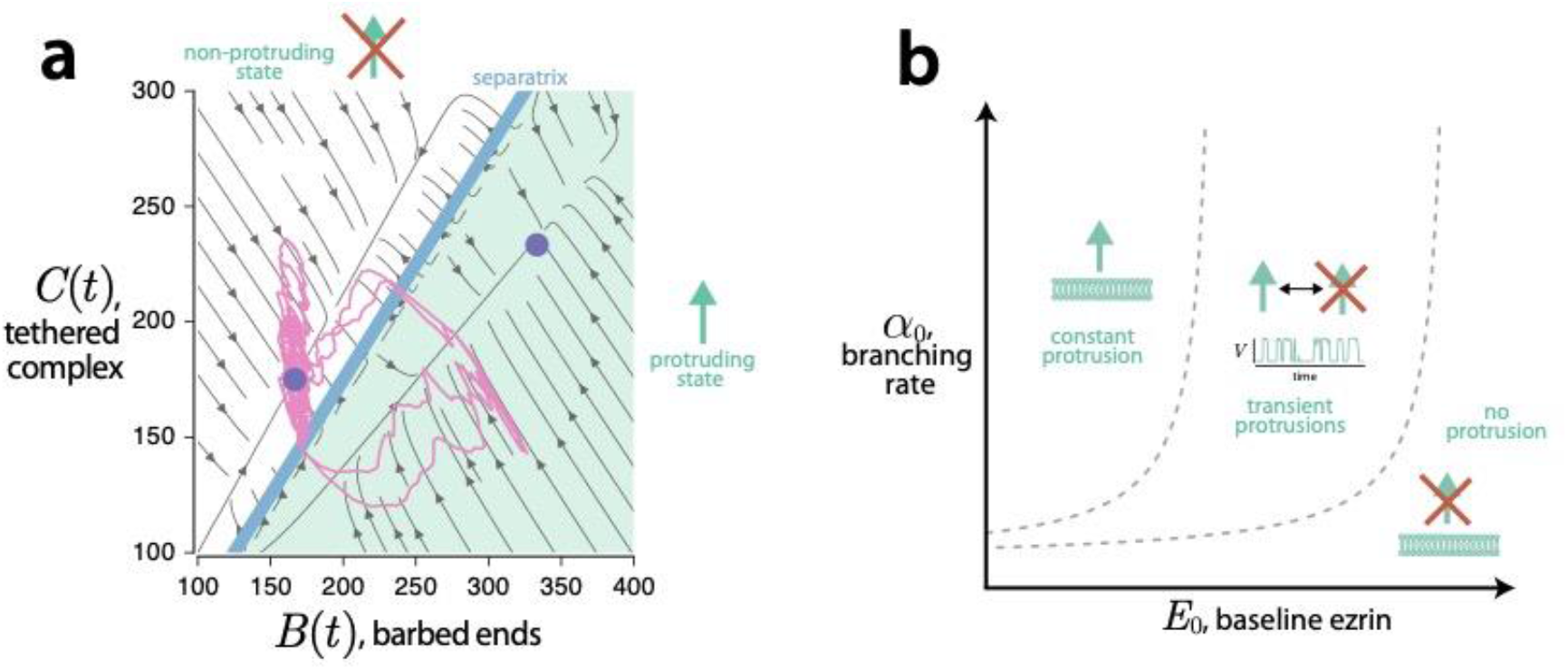
*On the left: a phase portrait of the typical simulation (with ezrin fluctuations) seen in the main text as a function of the number of barbed ends and complexes. Protrusive events can be interpreted as a metastable switch across the separatrix. The two stable equilibria are labeled in purple circles. On the right: the condition* (7) *as a function of the baseline ezrin level E*_0_ *and branching rate a*_0_.

With this analysis, we now understand protrusive events in the nonspatial model as metastable switching between the two equilibria across the separatrix (barrier), as seen in **Methods Figure 3a**. We note that the simulations use a finite *λ* and therefore have true equilibria that deviate slightly from the ones used for the analysis. Consequently, the simulation seen in the figure does not relax to exactly the predicted equilibria, but still demonstrates the qualitative behavior.

Furthermore, this analysis gives us insight toward the interplay of actin and ezrin levels, as seen in **Methods Figure 3b**. So long as the branching rate is sufficiently high to overcome membrane tension, ezrin levels primarily dictate the protrusive behavior. That is, moving vertically in the diagram (changing actin levels) produces little difference in regime in contrast to moving horizontally (changing ezrin levels).

#### Ezrin buildup model

We hypothesize that during retraction (or stall) events, the motion of the membrane pools ezrin in the cytoplasm or in other unavailable stores near the membrane. This results in a buildup of ezrin near or on the membrane, but unavailable for actin binding. When a protrusion event occurs, this pool dissipates. To describe this phenomenon, we introduce this second pool of ezrin, denoted *E*(t)* which undergoes the dynamics

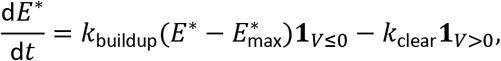

For simulations, we take 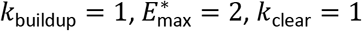, where *E*\* is again in arbitrary normalized units. The total ezrin is then

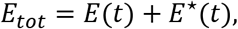

which is the quantity in the main text figure 2F. We note that this quantity, *E*\* has no feedback on the model and serves as a hypothesized explanation for the observed peak in ezrin prior to initiation of protrusive events.

#### Model experiments

In this section, we describe in more detail the experiments run on the mathematical model presented in the main text.

#### Actin vs. ezrin fluctuations

To explore how different sources of stochasticity influence protrusions, we slightly modify the model by removing ezrin fluctuations and adding actin fluctuations. This is of natural interest out of the possibility that just fluctuations in the number of barbed ends (or Arp2/3 levels) could drive protrusions. Specifically, in these simulations, we take *σ_E_* = 0 and change the dynamics of the barbed ends, *B*(*t*) to be

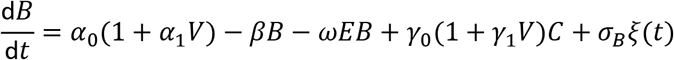

with *σ_B_* = 25. The result of these simulations, compared with the original model can be seen in main text figures 2C and 2D. From these, we find a critical conclusion: fluctuations in actin seemingly *cannot* drive protrusion events. The reason for this is as follows: If the ezrin pool is sufficient, then if the amount of barbed ends fluctuates larger than its baseline value, some fraction of these barbed ends will become tethered to ezrin, meaning the ratio of free to tethered barbed ends remains relatively unchanged, disallowing a protrusive event. However, if the free ezrin fluctuates, this ratio of free to tethered barbed ends may change significantly, allowing for protrusions. This is summarized graphically in **Methods Figure 4**.

**Methods Figure 4.**
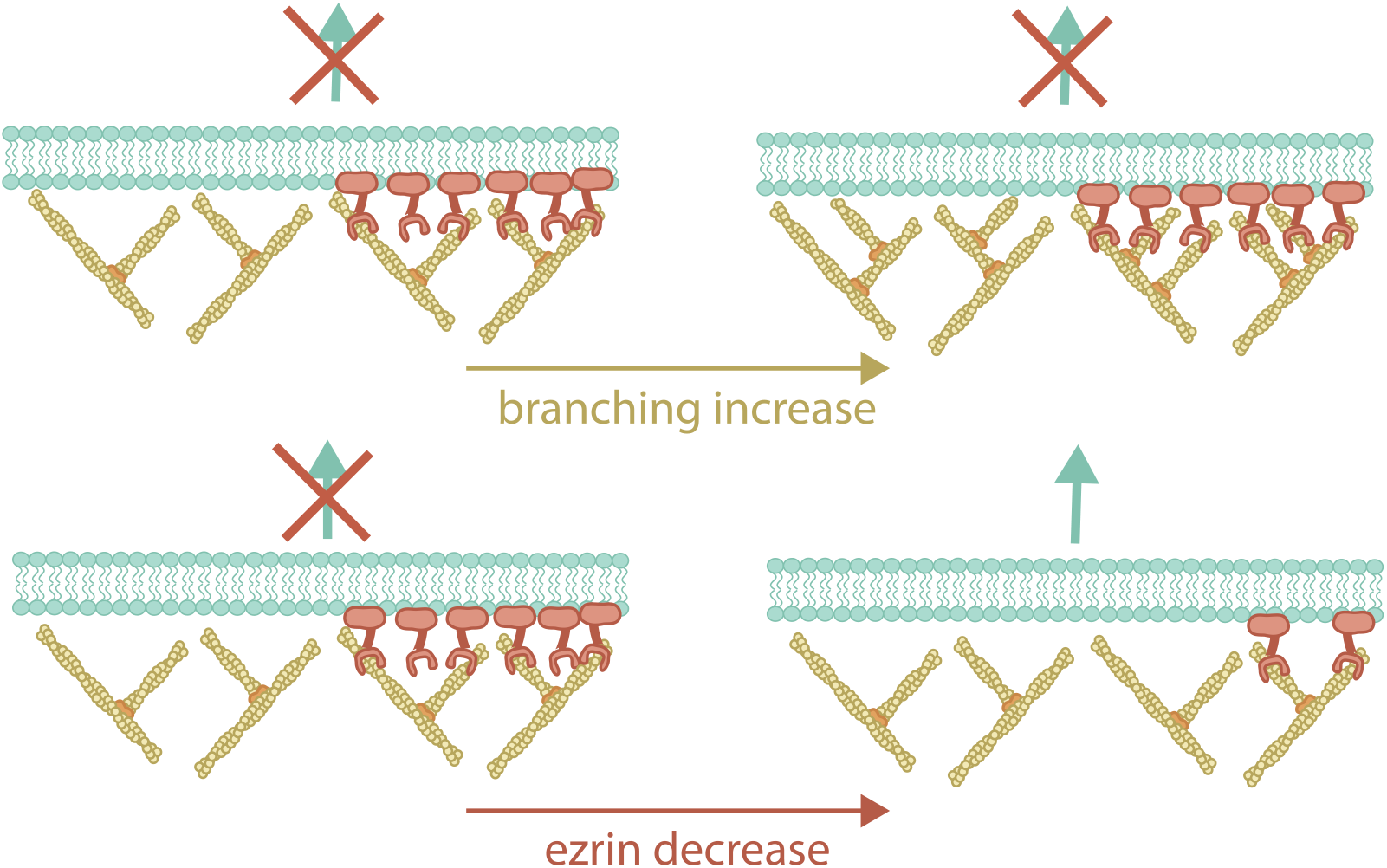
Cartoon explaining the fundamental difference between actin (or Arp2/3) fluctuations and ezrin fluctuations. Actin fluctuations do not alter the ratio of working to tethered, and therefore cannot drive protrusions. However, ezrin fluctuations do alter this ratio and can initiate protrusions.

#### Introduction of high-affinity ezrin

The introduction of ezrin TD was simulated in the model by introducing a separate pool of ezrin with higher affinity *(51)*, but structurally looks no different. Call this second population 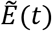. Then, the system (2) becomes

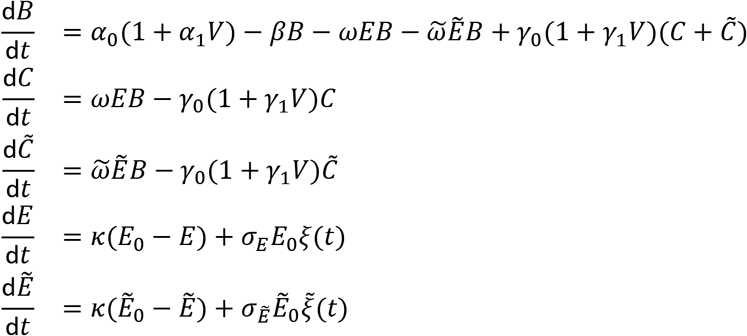

To explore how the relative quantities of regular and TD ezrin contribute, we take 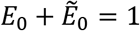. That is, we specify that some fraction *E*_0_ = *θ* ∈ [0,1] of the population is regular ezrin with affinity *ω* and then the remaining portion 1 – *θ* is TD ezrin, with high affinity 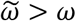. The results of sweeping over this fraction *θ* can be seen in main text figures 4G, 4J, 4M.

#### Varying Arp2/3 expression

Arp2/3 levels are not directly included in the model, but the F-actin nucleation/branching term is assumed to be Arp2/3 mediated, and therefore manifests in the model in that way. Thus, to simulate varying the Arp2/3 levels at the leading edge, *α*_0_, the branching rate varied from 50% of its original value (underexpressed) to 150% of its original value (overexpressed). Qualitatively, as Arp2/3 levels and therefore branching increases, there is an increase in protrusive activity, seen in figures 4M, 4L of the main text.

This seems like it contradicts the point that actin fluctuations cannot drive protrusions, but it is a distinct point. Increasing *α*_0_ increases the total amount of F-actin in the system, which does not initiate a protrusive event on its own. The velocity is determined by not only the ratio of the working and tethered filaments, but also the inherent membrane tension. Consequently, increasing the overall levels of actin does not affect the ratio of filaments but does help overcome the membrane tension, driving the system closer to a protrusion event overall.

#### Model sensitivity analysis

Sensitivity analysis was performed by normalizing the relative effect of a parameter change, measured in each of protrusion frequency, duration, and velocity, by the relative change in that parameter. The statistical comparison shown in Figure 4N is a t-test of the means of the model sensitivities calculated for the following parameter values that modulate arp2/3 activity and ezrin affinity. Simulated protrusion duration showed no significant difference due to changes in either arp2/3 activity or ezrin affinity (data not shown).

## Supplementary Figures

**Supplementary Figure 1.**
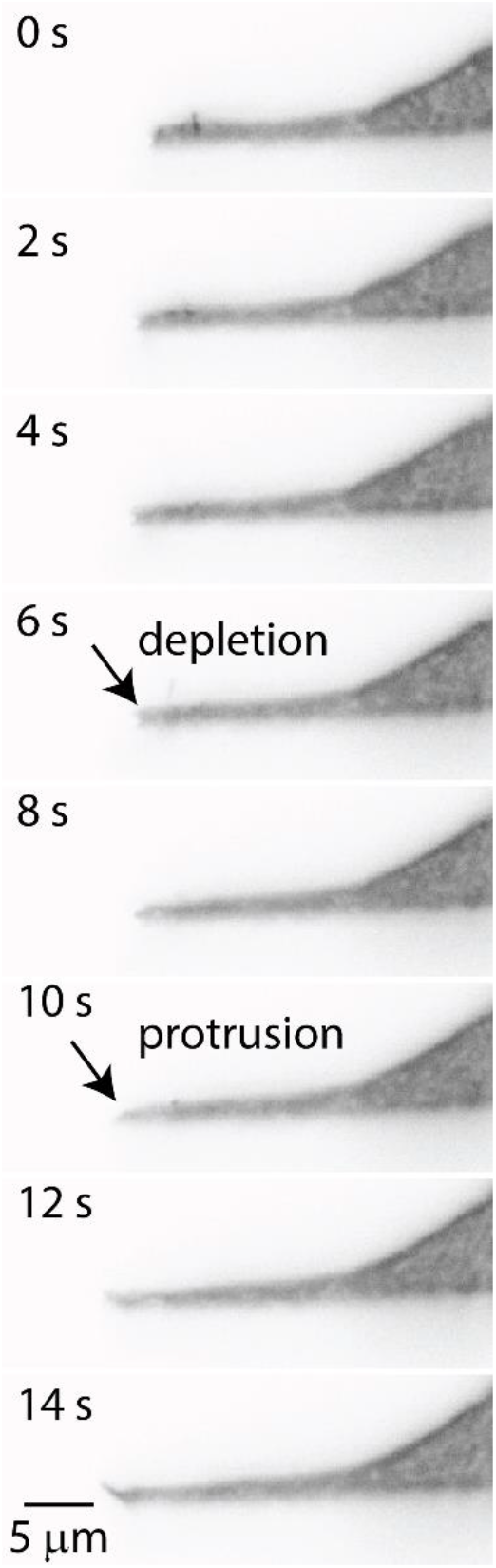
Lateral-axial (i.e., x-z) reslice of light sheet fluorescence image of GFP-ezrin in U2OS cells showing that within the diffraction limit, nascent protrusions do not appear thinner than other lamellipodial regions.

**Supplementary Figure 2.**
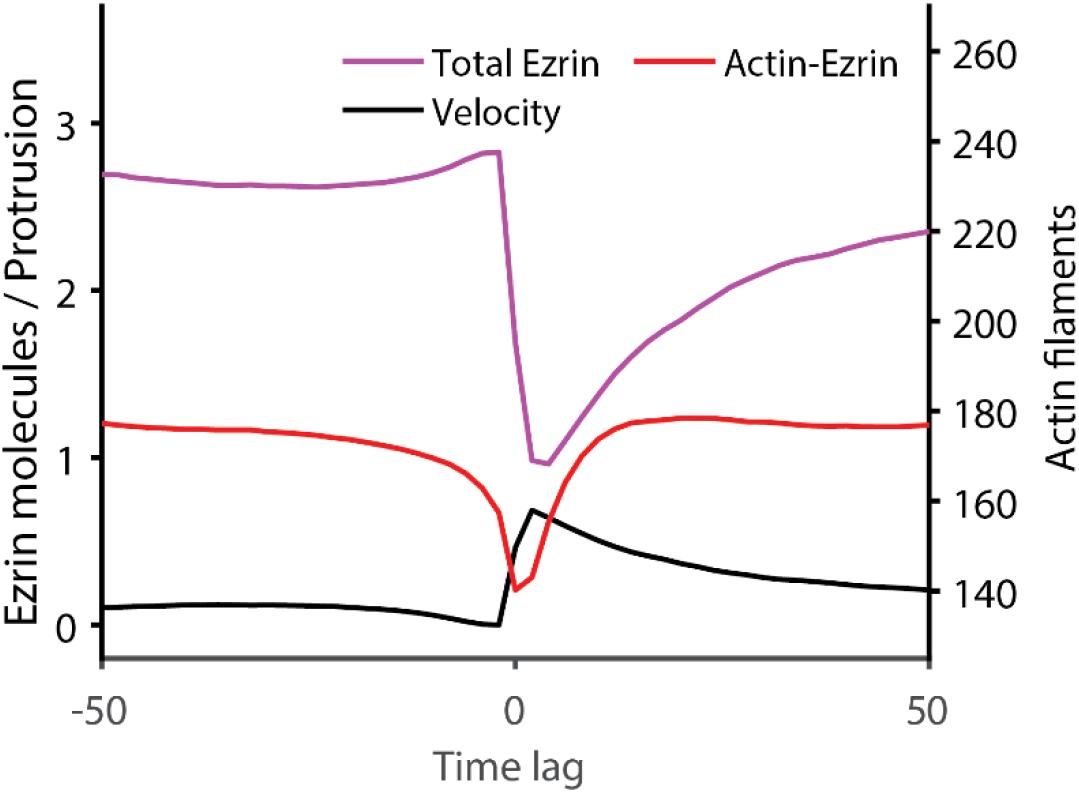
Ezrin accumulation model, showing how ezrin may build up in cytoplasmic stores during retraction. Average traces of total ezrin, F-actin-bound ezrin and edge velocity are aligned to protrusion onset.

**Supplementary Figure 3.**
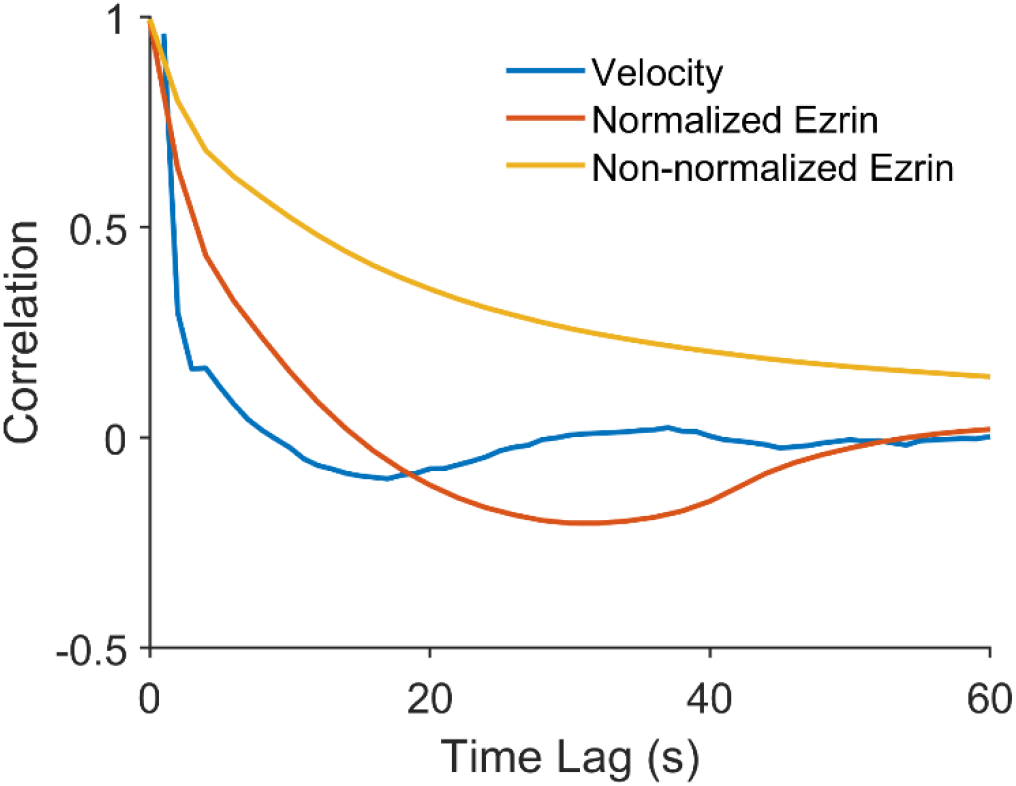
Autocorrelation function in velocity, ezrin, and low frequency normalized ezrin, showing how low frequency normalization focuses on ezrin fluctuations of the same time scale as protrusion fluctuations.

**Supplementary Figure 4.**
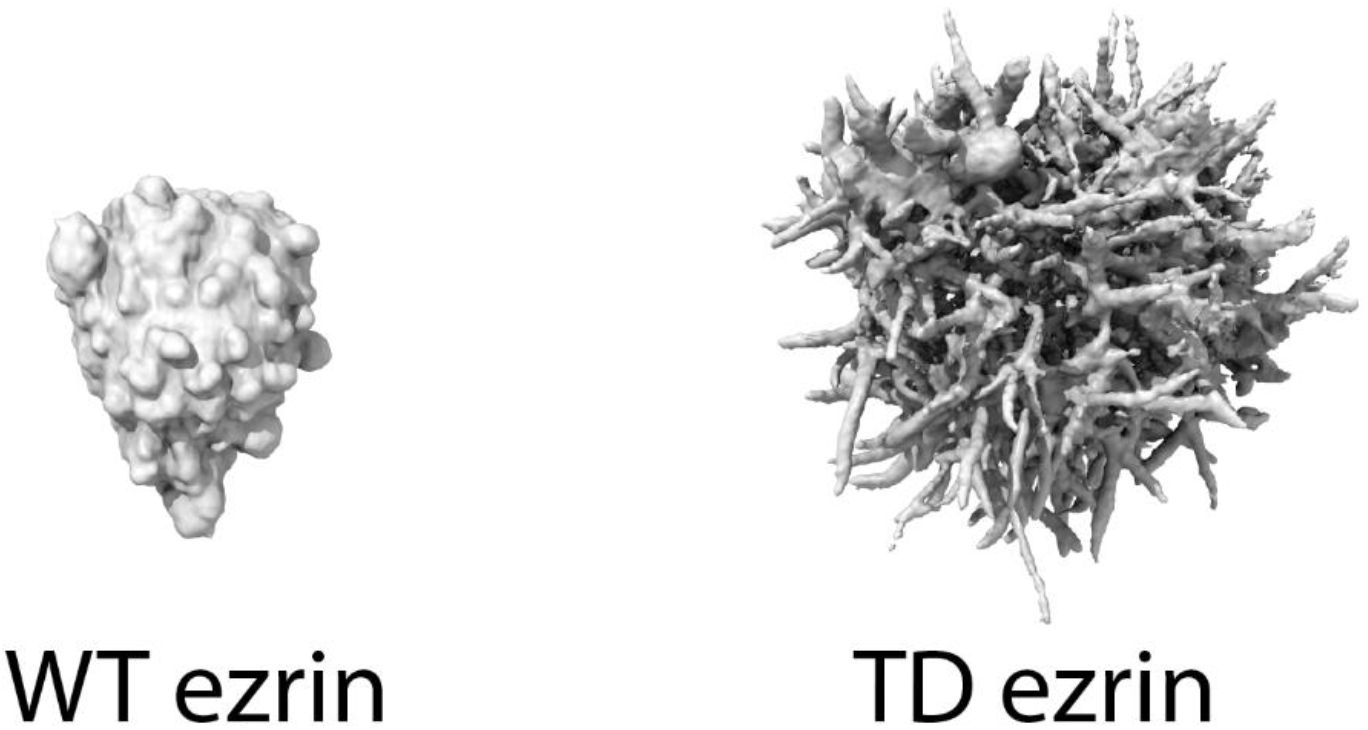
Surface of a melanoma cells expressing wildtype (WT) vs T567D (TD) mutant ezrin, imaged in 3D using light sheet microscopy.

**Supplementary Figure 5.**
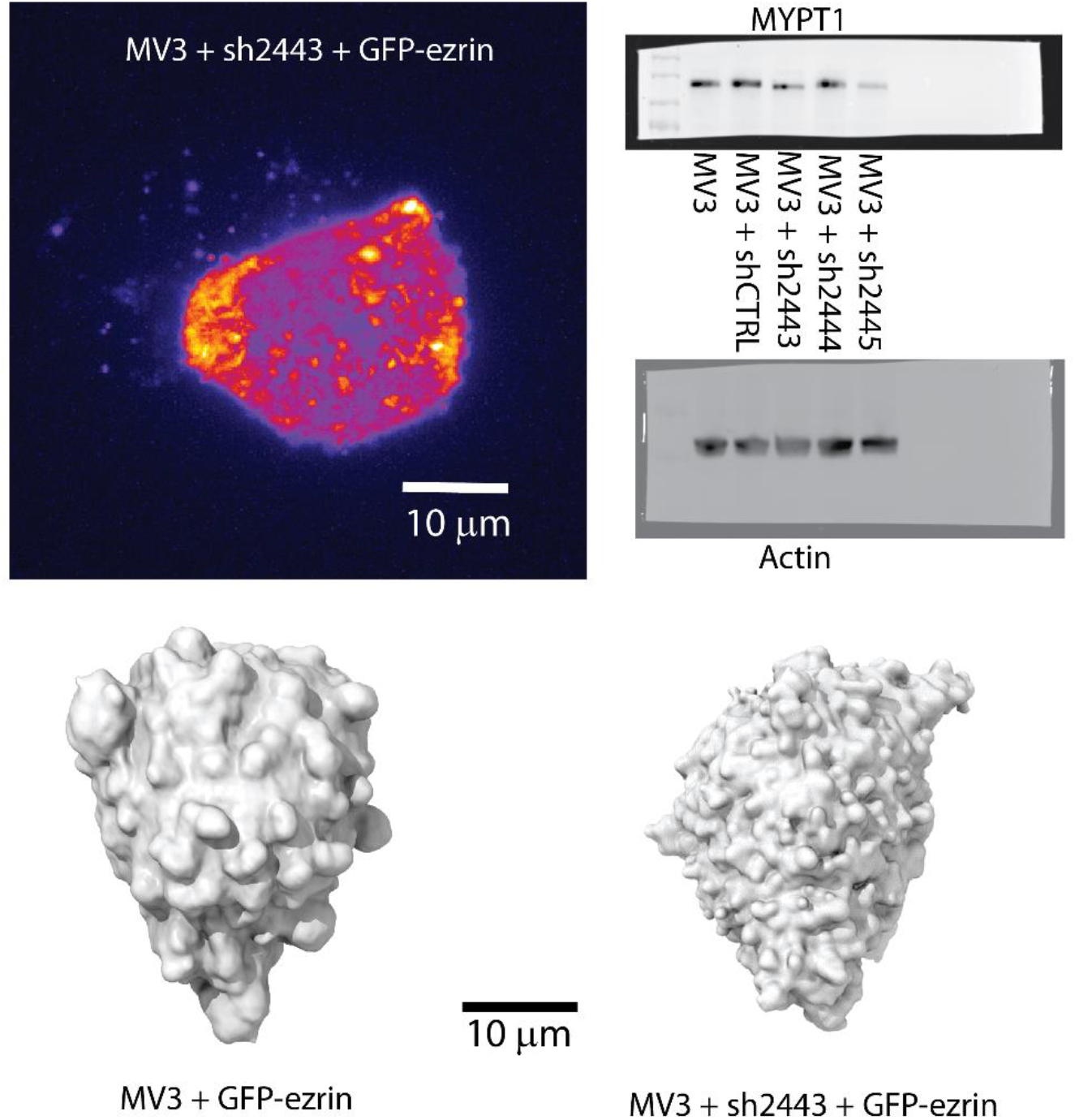
Maximum intensity projection of a melanoma cell expressing GFP-ezrin, under knockdown of ezrin phosphatase MYPT1, imaged in 3D collagen by light sheet microscopy. Western blot confirming knock-down of the phosphatase using three shRNA oligonucleotides. 3D surface renderings of melanoma cells without and with MYPT1 knockdown.

**Supplementary Figure 6.**
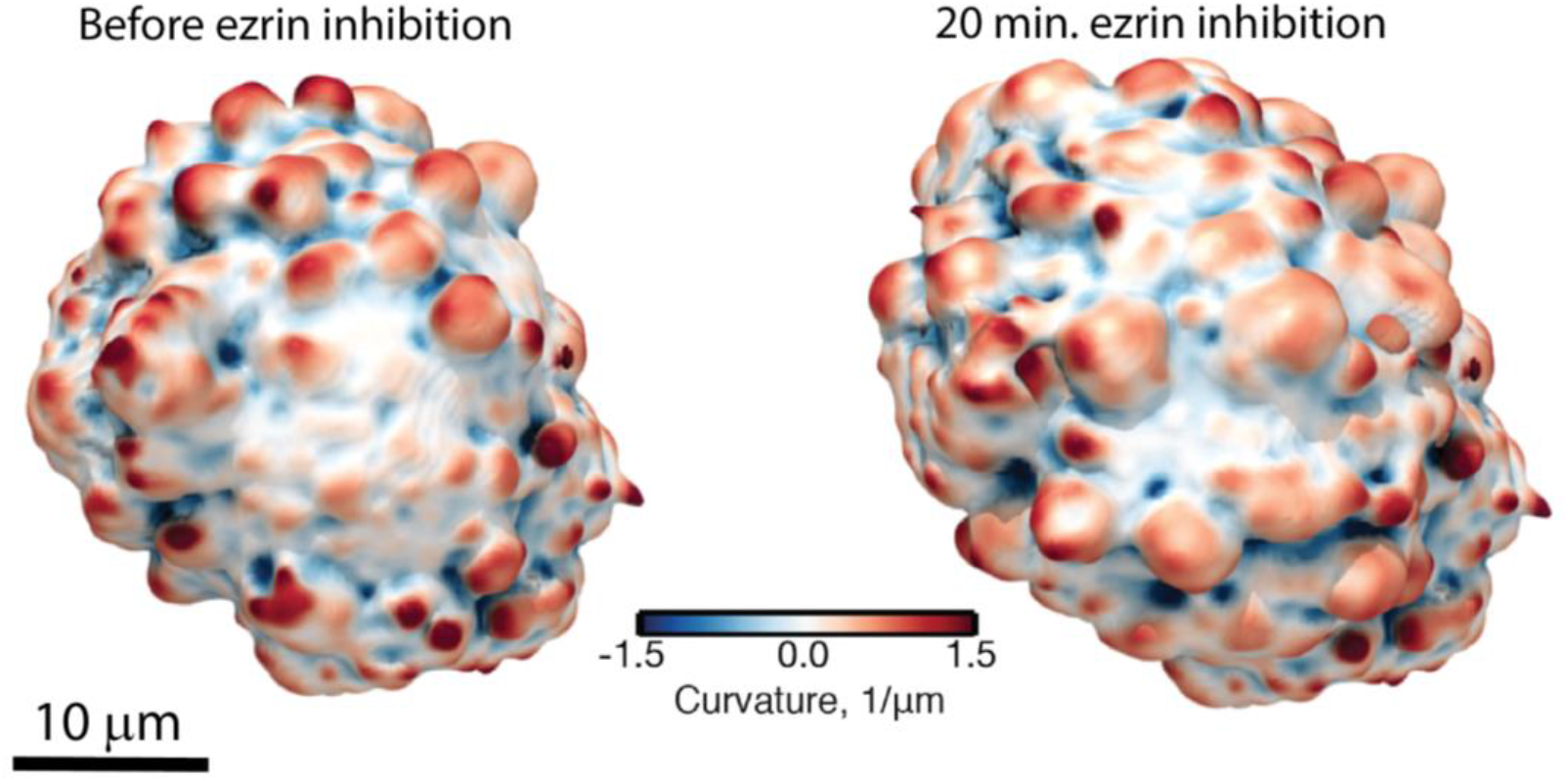
Renderings of the surface curvature of a melanoma cell before and after ezrin inhibition via 10 μM NSC668394, imaged in 3D collagen using light sheet microscopy.

**Supplementary Figure 7.**
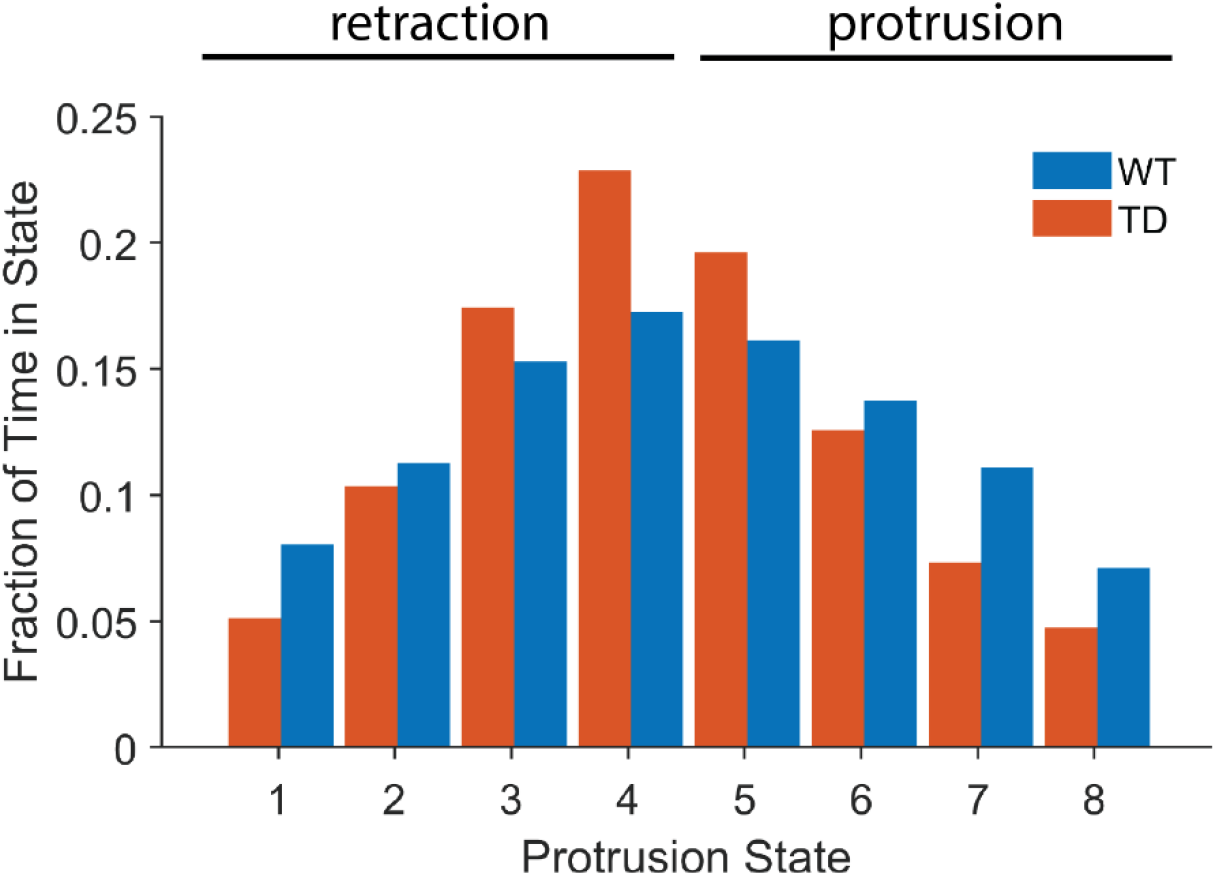
Fraction of time spent in different HMM-classified protrusion states in cells expressing ezrin WT or ezrin TD.

**Supplementary Figure 8.**
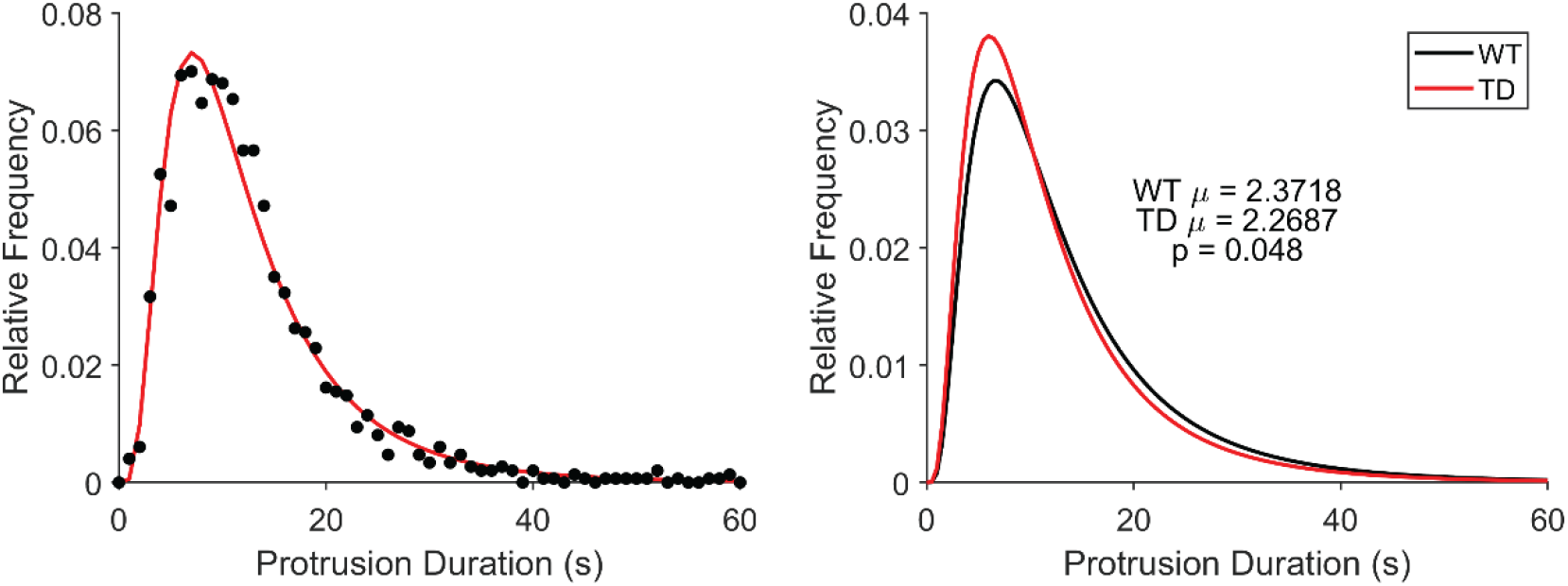
Fit of measured protrusion duration data to a lognormal model. P value.

**Supplementary Figure 9.**
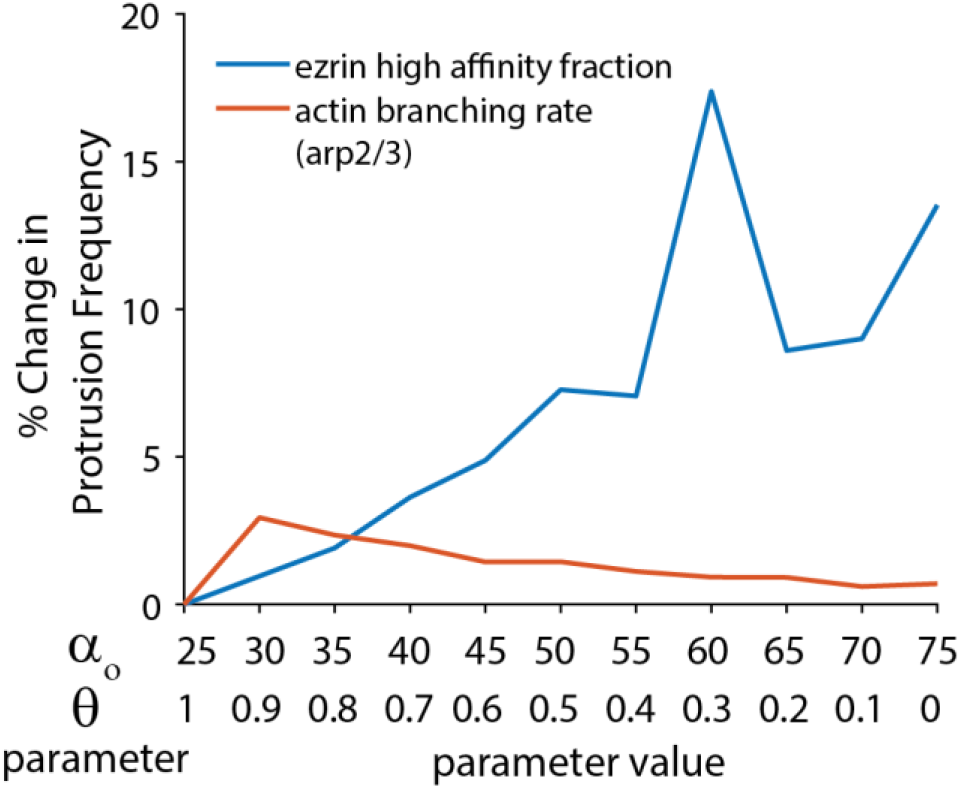
Relative change in protrusion frequency in model simulations as a function of the different parameter values listed in the Model Sensitivity Analysis subsection of the Materials and Methods. **add values and means lines.

**Supplementary Figure 10.**
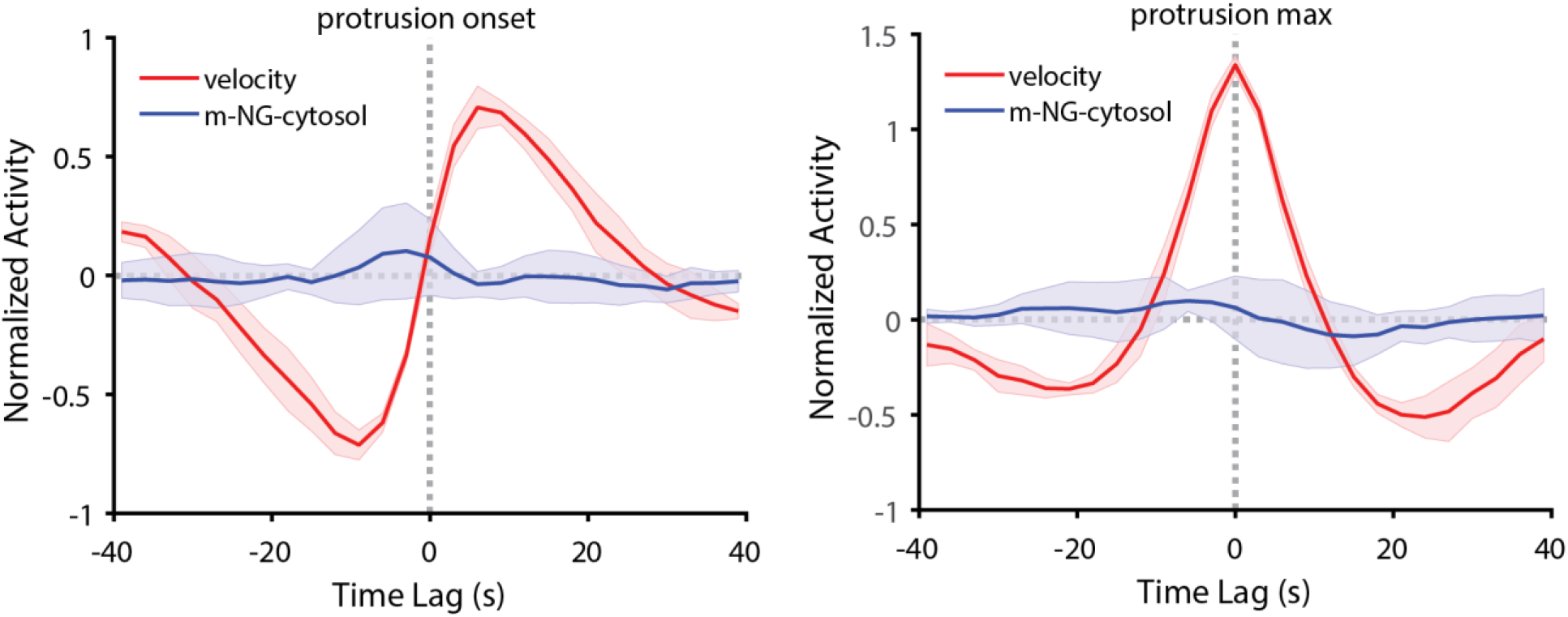
Analysis of cytosolic fluorescence relative to protrusion onset and protrusion maximum in U2OS cells.

**Supplementary Figure 11.**
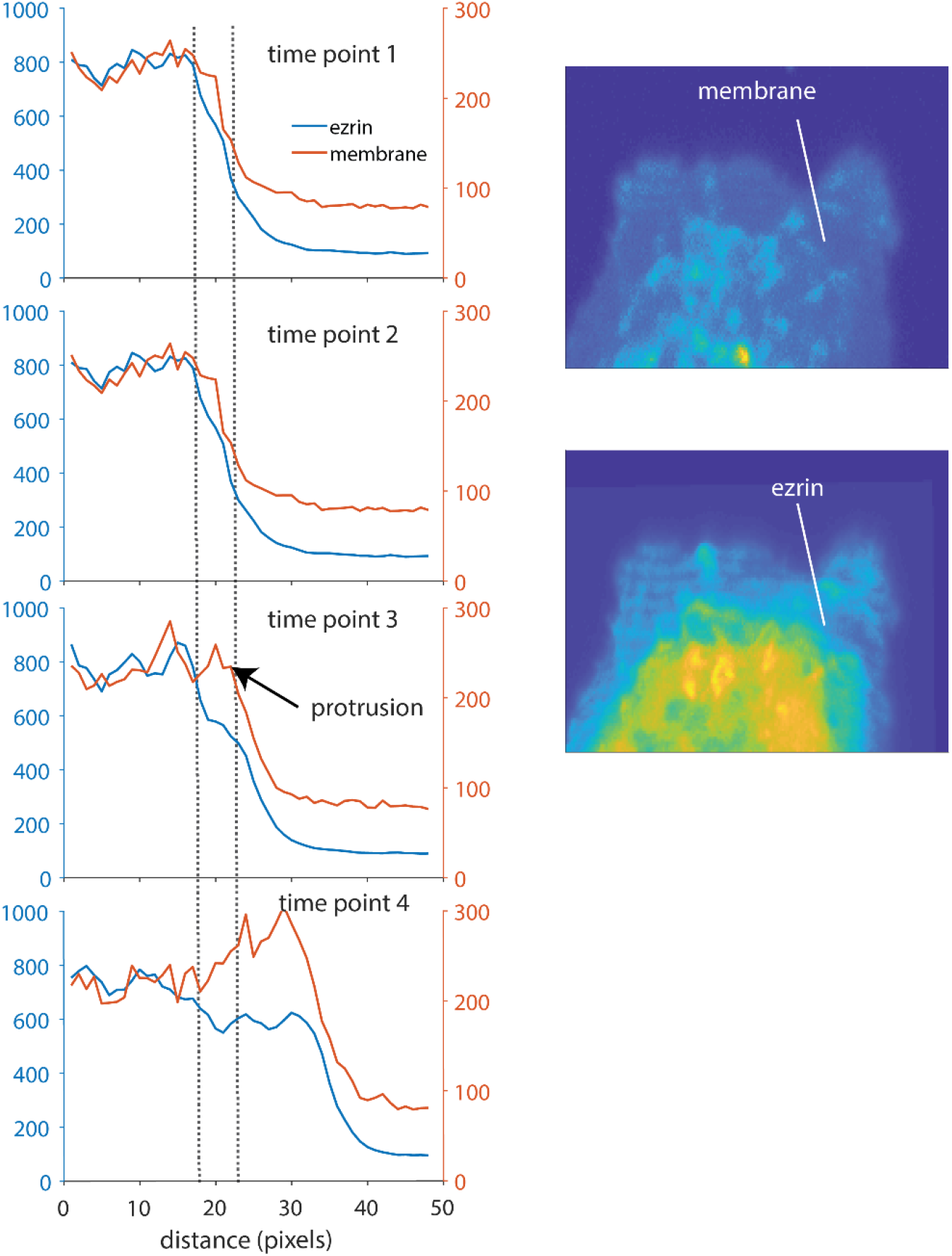
Linescan analysis of GFP-ezrin and membrane marker in U2OS cell, showing that membrane protrusion precedes a recovery in GFP-ezrin following protrusion. **Scale bars.

## Movie Legends

Supplementary Movie 1. Animation showing collagen surrounding MV3 melanoma cell.

Supplementary Movie 2. Cell surface motion of an MV3 melanoma cell. Colors indicate protrusion (purple) and retraction (green) as indicated in the colorbar in Figure 1B.

Supplementary Movie 3. GFP-ezrin intensity projected onto the surface of an MV3 melanoma cell. Colors indicate high (red) and low (blue) ezrin concentration as indicated in the colorbar in Figure 1C.

Supplementary Movie 4. Maximum intensity projection of GFP-ezrin in amoeboid MV3 melanoma cell. Colors indicate ezrin concentration on a non-linear and saturated scale as indicated in the colorbar in Figure 1D.

Supplementary Movie 5. Maximum intensity projection of GFP-ezrin in mesenchymal MV3 melanoma cell. Colors indicate ezrin concentration on a non-linear and saturated scale as indicated in the colorbar in Figure 1D.

Supplementary Movie 6. Maximum intensity projection of GFP-ezrin in a neuron. Colors indicate ezrin concentration on a non-linear and saturated scale as indicated in the colorbar in Figure 1D.

Supplementary Movie 7. Maximum intensity projection of GFP-ezrin in a U2OS cell, imaged using light sheet microscopy. Colors indicate ezrin concentration on a non-linear and saturated scale as indicated in the colorbar in Figure 1D.

Supplementary Movie 8. GFP-ezrin in a U2OS cell, imaged using spinning disk microscopy. Colors indicate high (yellow) and low (purple) ezrin concentration as indicated in the colorbar in Figure 3A.

